# Stable coexistence and transport of lytic phage infections with migrating bacterial hosts

**DOI:** 10.1101/2025.04.18.649568

**Authors:** Jacopo Marchi, Chadiga Khalek, Abir George, Joshua S. Weitz, Remy Chait

## Abstract

Bacteriophages (phages), viruses that exclusively infect bacteria, coexist with their bacterial hosts across diverse environments at densities exceeding 10^7^ ml^-1^ in marine surface waters, 10^8^ ml^-1^ in soils, and 10^9^ ml^-1^ in the human gut^1–6^. In contrast, phage lysis of bacteria populations within well-mixed *in vitro* environments select for the emergence of phage-resistant bacterial mutants^7^, which in turn select for host-range expansion phage mutants^8,9^ and, typically, the collapse of phage populations altogether^10,11^. This gap in outcome raises a question: what enables long-term phage-bacteria coexistence? Here, we show how interactions in space can facilitate stable coexistence and long-range transport of virulent phage along with migrating bacteria. Combining theory, simulation, and experiments across multiple phage-bacteria systems, we reveal a chemotaxis-driven mechanism that robustly stabilizes coexistence and dispersal of virulent phages with migrating hosts. These findings suggest the ecological relevance of spatial interaction mechanisms that reinforce stability between antagonistic partners in the absence of perpetual cycles of defense and counter-defense, which may be broadly applicable across phage-bacteria systems.

## Introduction

Phage infection can lead to the rapid takeover of cellular machinery, phage genome replication, self-assembly of capsids, cellular lysis, and the release of hundreds of infectious virions^12–15^. The death of individual cells does not necessarily eliminate bacterial populations^16,17^. Instead, obligately lytic phage and their bacterial hosts can coexist over multiple replication cycles given predator-prey like dynamics as predicted via nonlinear dynamic theory^16^ and validated in chemostats^18–20^. On longer time scales, strong selection to resist phage-induced lysis by bacteria and corresponding pressure to infect mutant bacteria by phage can drive complex eco-evolutionary processes that include arms races and Kill-the-Winner dynamics^10,21,22^. Yet, efforts to recapitulate persistent coexistence between lytic phage and bacteria in the laboratory are often limited. Coevolution in well-mixed laboratory settings typically leads to the emergence of phage-resistant bacterial mutants that phage cannot infect even after multiple rounds of coevolution^10,11,23^, consistent with a hypothesized ‘asymmetric advantage’^8^ in bacterial evolutionary potential relative to phage.

In contrast to well-mixed laboratory conditions, population dynamics and community structure of complex microbiomes are shaped, in part, by spatially explicit interactions^24–27^. Spatially-structured systems can exhibit markedly different behaviour than their well-mixed counterparts – including the potential for sustained diversification and coexistence amongst parasites and hosts^25,27–32^. Spatial structure can also reshape phage-bacteria coexistence. Motile bacteria can migrate orders of magnitude faster over ecologically relevant timescales than is possible for phage whose mobility is driven by diffusion. While phage can infect and proliferate quickly, bacteria that move beyond zones of infection can escape infections. The potential for motility-enabled escape from parasitism is analogous to theories underlying the evolution of long-distance seed dispersal in plants^33,34^ and raises questions on whether spatial interactions enhance or suppress phage-bacteria coexistence.

The impact of spatial interactions on phage-bacteria coexistence is particularly relevant for bacteria that migrate via chemotaxis^35^. In well-studied model systems, including *Escherichia coli*, the local growth of cells depletes resources and induces chemical gradients such that chemotactic bacteria swim towards resource-rich environments (and associated chemoattractants). The feedback between chemical uptake and cellular movement leads to emergent, collective migration in the form of range expansion waves with sharply defined population fronts^36,37^. Multiple studies have used chemotactic bacteria as a lens to examine whether and how phage can coexist – and even keep up – with motile bacterial populations^38–40^. In doing so, these studies have explored (i) alternative population patterns when a migrating bacterial front encounters a localized inoculum of phage^38^; (ii) the relevance of phage-resistance on transport-mediated coexistence when phage and chemotactic bacteria are coinoculated together^39,40^. Combining mathematical models and experiments together, these studies have shown that phage can propagate with hosts in localized clearance domains over short length scales on the order of a few centimeters within agar plates, but that such patterns may be unstable as small perturbations lead to alternative outcomes, i.e, spreading phage lysis of bacteria or bacterial escape^38^. In addition, phage propagation may be sensitive to the emergence of phage resistance amongst hosts^39^ or may be locally limited insofar as phage-resistant bacterial mutants heterogeneously colonize previously lysed domains^40^. Together, previous studies raise a key question: can phage robustly and stably co-propagate with chemotactic bacteria over longer spatiotemporal scales?

Here, we used timelapse imaging of agar swim assays to address the impact of spatial interactions on coexistence of virulent phage with chemotactic *E. coli* and *Pseudomonas aeruginosa* strains. Our large format agar plates enable us to explore the long-term emergent dynamics of migrating bacterial hosts that encounter virulent phage in spatially explicit contexts. As we show, virulent phage can propagate with migrating bacterial fronts at speeds an order of magnitude faster than through the non-migrating bacterial lawns behind the colony expansion front. Notably, phage infection transport remains stable over multiple days and across distances exceeding tens of centimeters. Through a combination of theory, simulation, and experiments, we identify quantitative principles underlying the emergence of stable infection transport in multiple phage-bacteria systems. We then characterize the stability and robustness of motility-mediated infection transport to environmental and ecological perturbations, showing how phage can dynamically coexist with chemotactic bacteria even when abrupt changes in collective cell migration would seem fast enough to leave phage behind.

## Results

### Lytic phage invasion and propagation with migrating bacterial fronts

We initiated spatial infection experiments by inoculating a localized 1µl drop of virulent phage lysate approximately 2 cm away from a phage-sensitive, chemotactic bacterial culture in large format, soft agar swim plates. By collecting high-resolution timelapse images of the expanding bacterial population and lytic clearing zones with a scanner-based platform (see Methods), we followed propagating host-phage dynamics for days over distances up to 25cm. Figure 1a shows the results of inoculating an LB swim plate containing 0.25% agar with virulent *Escherichia phage* T7 and *E. coli* K12 strain W1485(F8) pre-adapted to swim assays to reduce the confounding emergence of “fast” mutants (for all strains see Methods). Phage T7 had a dramatic effect upon encounter by the migrating bacterial front, rapidly lysing hosts and inducing the emergence of a widening front invasion clearance zone. This clearance zone was then protected from immediate bacterial re-invasion (Figure 1A, 14-21h, Supplementary Movie 1, and Supplementary Figure 1). Despite the dramatic lysis-induced disruption of the expanding bacterial front, the chemotactic bacterial population continued to propagate outward without interruption in regions outside the clearance zone, slightly outpacing the infection and curving gradually inwards (Figure 1A, 21h-27h). As the two propagating bacterial fronts on either side moved inward, the phage cleared zone grew narrower and the inward curvature ceased (Figure 1A, 31h). Once the two fronts converged, the phage infection coalesced to a focused clearance zone and then propagated stably with the bacterial front at a speed of ∼5mm/hr (Figure 1A, 31h-48h), an order of magnitude faster than T7 plaque expansion in static bacterial populations^41^. Note that the early, approximately straight radial lysis boundary of the invasion zone aligns with previous observations of short-term invasion and co-propagation of phage χ with migrating *E. coli* and *S*.*typhimurium* at the scale of 9 cm diameter plates^38^ (dashed line in Figure 1A).

**Figure 1.**
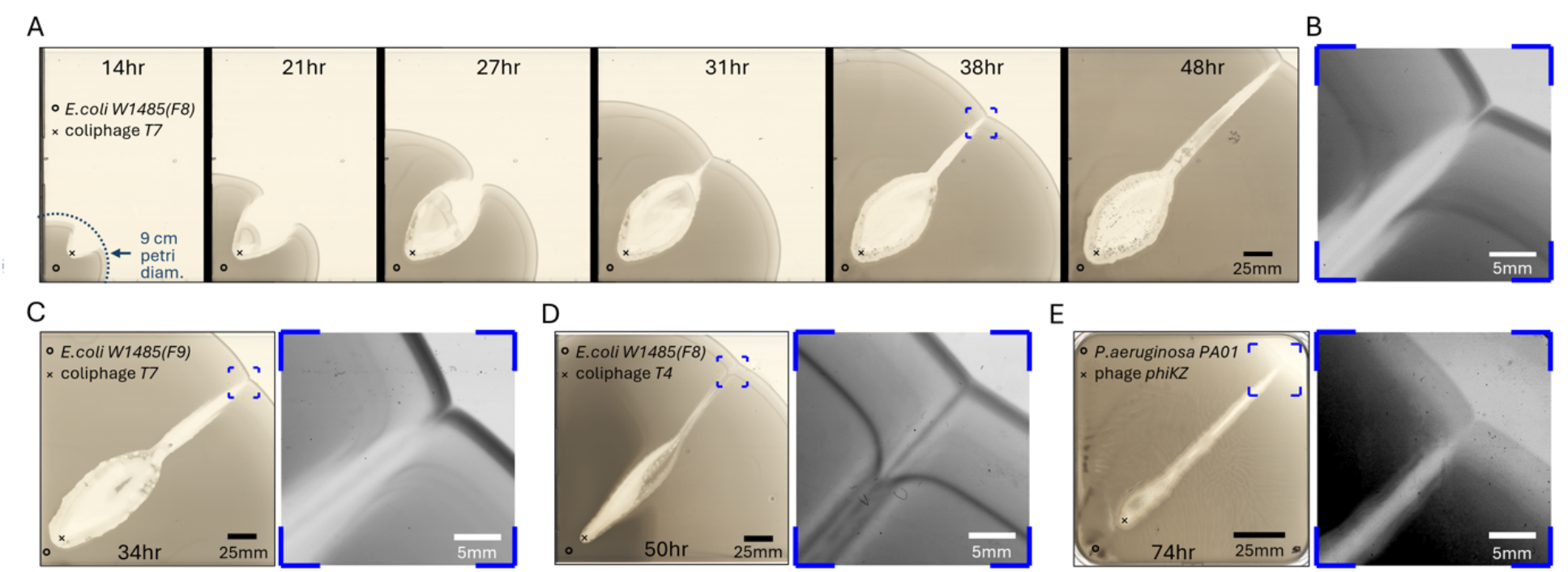
Extended regions cleared by lytic phage emerge and coalesce to stably propagating infections on chemotactically migrating bacterial fronts. **(A)** Time-lapse transmission scans of an expanding *E. coli* W1485(F8) population, moving chemotactically through 0.25% agar nutrient media from point “o” near the corner of a 24cm plate. The bacterial front is invaded by virulent T7 phage particles it encounters at point “x”. While still diverging at the scale of a standard, center-inoculated 9 cm swim assay (dashed arc) after 14hr, the phage-cleared region reconverges by 30hr and settles into a narrow stable propagation across the rest of the test plate. Scale bar 25mm. **(B)** A zoomed, linearly contrast-enhanced view of the stably propagating infection front in panel A at 38hr (blue box). Similar virulent phage invasions, followed by expansion, reconvergence and stable extension of infections with migrating host fronts is seen for **(C)** W1485(F9), an *E. coli* W1485(F8) mutant with a single dominant front, encountering phage T7, **(D)** *E. coli* W1485(F8) encountering phage T4, and **(E)** *P. aeruginosa* PA01 encountering phage ΦKZ. Transmission scans (left) and linearly contrast-enhanced zooms (right) are shown for indicated 25mm regions (blue boxes).

We repeated the experiment with *E. coli* K12 strain W1485(F9), a mutant of W1485(F8) with a single, dominant chemotactic front which migrates approximately 40% faster (∼7.2mm/hr) in the same conditions. Encounter of phage T7 by this faster single-front *E. coli* mutant leads to qualitatively similar dynamics, including a transient front invasion clearance zone, inward curvature of bacterial fronts on either side of the clearance zone, and coalescence to a stably propagating infection and phage-cleared zone in the center of the migrating bacterial front (Figure 1C). Stable infection transport also emerged across other phage and host bacteria combinations we tested, *e*.*g*., phage T4 and *E. coli* W1485(F8) (Figure 1D) and jumbo phage phiKZ and *P. aeruginosa* PA01 (Figure 1E). In each case we observed the stable propagation of narrow phage-induced clearance zones originating at the leading edge of the migrating bacterial front (see blue highlighted panels in Figure 1B-E). Stable infection transport dynamics similarly emerge for the ancestral W1485 and widely-studied MG1655 strains of *E. coli* (see Supplementary Figure 2). We refer to the propagating phage infection and associated cleared zone within bacterial chemotactic fronts as a Phage-Amplified Spatially Extended Region or “PhASER”. The stable establishment and persistence of PhASERs across multiple phage-bacteria systems with distinct bacterial chemotactic mechanisms and phage life history traits suggests the relevance of common, eco-physical principles.

### Stabilization of phage transport along population-mediated chemical gradients

To explore the basis for PhASER emergence, we developed a nonlinear, partial differential equation (PDE) model of the spatiotemporal dynamics of phage-sensitive and infected bacteria, virulent phage, resources, and chemoattractant molecules (see Box 1). The model combines quantitative principles of chemotactic range expansion in bacteria^36^, with infection-mediated feedback mechanisms used in ecological models of phage and bacteria^17,41^. In this reaction-diffusion-advection model, bacteria take up resources, reproduce, and migrate up gradients generated by chemoattractant depletion while phage adsorb to and infect bacteria that themselves continue to migrate, eventually lysing and releasing infectious virions back into the soft agar environment (see Box 1 for a schematic and overview of model ingredients and Supplementary Text for a complete description of model equations and parameters). The underlying reaction-diffusion-advection model includes key feedback terms, proposed in foundational models of phage dynamics: (i) phage lysis rate and burst size are inhibited in resource-depleted environments, decreasing with reductions in bacteria growth rate^42,43^; (ii) phage infections are explicitly structured in multiple stages to account for unimodal latent period distributions^41,44^. These elements were included in some^39^ but not all^38^ prior spatial models of phage infections of chemotactic bacteria. In contrast to these previous studies, the PDE approach we propose includes a Hill function modulating phage adsorption to account for discrete effects and avoid phage invasion from infinitesimally small densities (see analysis in the Supplementary Text, Section 3) and incorporates a generalized nonlinear phage adsorption profile saturating at high phage concentrations, following observations of phenomenological Michaelis-Menten reaction kinetics^45,46^. Finally, we excluded bacterial and phage evolution, to focus on emergent stable co-propagation of phage infections and hosts rather than resistance regrowth into previously lysed regions (as experimentally reported in ^40^).

The resulting reaction-diffusion-advection model generates stable phage invasion and transport patterns at population scales over long distances and time scales that are both qualitatively and quantitatively consistent with PhASERs observed in large-format agar plate experiments. Figure 2 shows model-simulated dynamics of phage infection and long-range transport using parameters estimated for *E. coli* W1485(F9) and phages T7 and T4 (see Supplementary Text) alongside experimentally measured data for the same strains (Supplementary Movies 2-5). In both cases, the model exhibits an initial expansion of a lytic clearance zone, coalescence to a focused infection in the chemotactic front, and the stable migration of this phage infection at the same speed as the migrating bacterial population. Curved lines overlaid over late-stage images (Methods, Supplementary Figure 3) show the correspondence of experimental and simulated migrating bacterial front positions at 2-hr intervals. Model simulations also exhibit qualitatively similar front invasion transients with wider phage T7 than T4 clearance zones before coalescence to a stable PhASER, and more rapid erosion of the static host lawn behind the fronts by phage T7.

**Figure 2.**
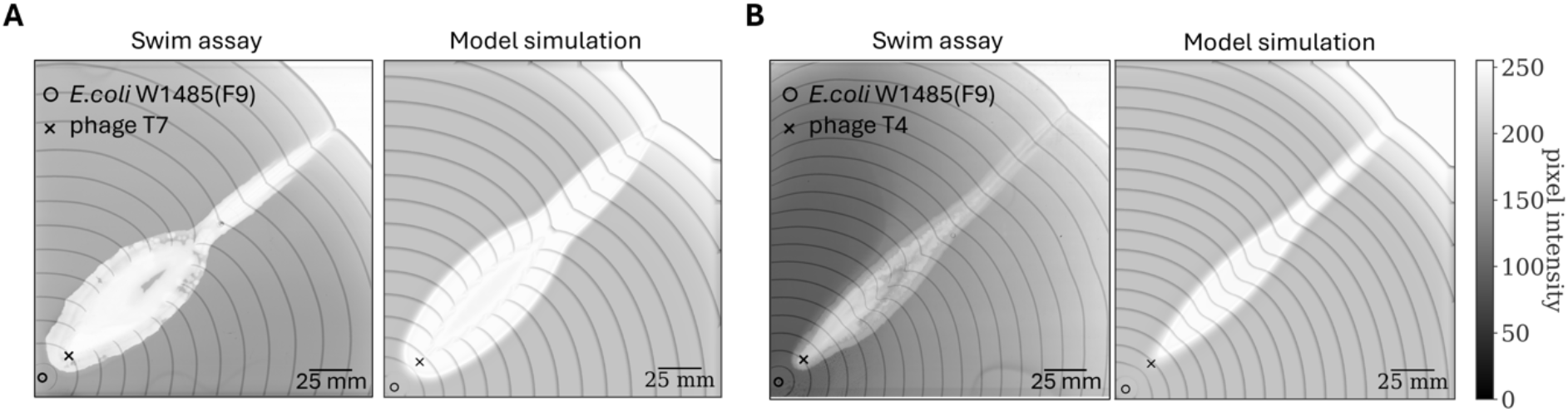
Phage Amplified Spatially Extended Regions (PhASERs) observed in experiments and model simulations. Images of populations (grey) of *E. coli* strain W1485(F9) growing and moving chemotactically from location ○, that encountered **(A)** T7 or **(B)** T4 phages at location × in 24x24cm soft agar swim assays. In both cases, elongated, clear invasion transients appear and resolve to narrow, stably propagating infections. Experimental data (blue channel of color transmission scans) are displayed to the left and model simulations (cell densities mapped to calibrated blue channel pixel intensities, Methods) shown to the right. The positions of the dominant chemotactic front (grey curves) of W1485(F9) are overlaid on each image at 2-hour intervals to highlight its shape and trajectory throughout the infection without confounding by continued phage lysis behind the migrating fronts (see also Supplementary Movies 2-5). All simulation parameters are indicated in Supplementary Text Table S1, except for A) where we modulate the bacteria diffusion constant *D*_*B*_ = 2.7 · 10^5^*μm*^2^/hrs and responsiveness to sensed gradient *χ* = 5.5 · 10^6^*μm*^2^/hrs to match the faster range expansion (around 7.2 mm/hr) in this experimental replicate.

We hypothesized that the emergence of PhASERs in the mathematical model depends on the inclusion of key mechanistic elements distinct from assumptions in prior, related work^38^. First, removing the resource dependency of infection and lysis leads to rapid, lateral expansion of phage clearing behind the front in contrast to the experimentally observed asymmetry between rapid infection front transport and slow lateral lysis expansion (Supplementary Figure 4 B & Supplementary Text). Second, if phage latent periods are assumed to be exponentially distributed, then the simulated lysis zone rapidly expands outwards near the front and continue diverging radially over long distances, contrasting with the reconvergence and PhASER formation that we observe in experiments (Supplementary Figure 4 A & Supplementary Figure 5). Next, reducing the complexity of the simulated growth medium by merging nutrients and attractants into a single state variable leads to qualitatively different infection propagation dynamics to those observed experimentally, with lysis boundaries appearing straight at any given time snapshot (Supplementary Figure 6). Finally, the robustness of experimental emergence of PhASERs with different phages (Figure 1) prompted us to assess the sensitivity of modelled dynamics to variation in key phage infection parameters. Our simulations show that PhASERs, while not inevitable, are robust to dramatic changes in phage characteristics including over 10-fold variation in adsorption rate and latent period (Supplementary Figures 7, 8 and 9). PhASERs are also robust to empirically-informed nonlinearities in the reduction of lysis rate and burst size with host growth rate^47^ (Supplementary Figures 10 and 11 & Supplementary Text).

#### Box 1

**Reaction-diffusion-advection model of phage infections of chemotactic bacteria**

<u>Summary:</u> We developed a nonlinear system of partial differential equations (PDEs) governing the reaction-diffusion-advection dynamics of chemotaxing susceptible (*S*) and infected (*E*) bacteria, phage *P*, resources *R* (magenta gradient) and chemoattractant *a* (green gradient). All population types diffuse according to a corresponding type-specific diffusion constant *D*. For bacteria, the diffusion represents the random component of a run-and-tumble process^48^ rather than Brownian motion. For viruses, resources, and chemoattractants, the diffusion represents that expected via Brownian motion.

<u>Reaction terms – bacterial uptake and growth:</u> All bacteria consume carbon, nutrients, and other small molecules in the complex LB broth – we refer to the compounds contributing to bacterial growth as resources, represented in cell density units (CFU/mL). Susceptible cells divide given a resource-dependent function 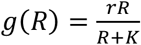, where *r* is the maximum growth rate and *K* is the Monod constant. We assume that resources consumed by infected cells are diverted to phage production, therefore infected cells do not divide. Bacteria also consume a separate set of molecules, the chemoattractants, according to a chemical-specific Monod law with rate 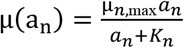. These are the chemical cues driving chemotaxis (see below) and are in principle distinct components to the resources (the main chemoattractants in the LB broth are aspartate and serine, see ^36^).

<u>Reaction terms – phage infection, lysis, and decay:</u> Phage adsorb to all bacteria and initiate infections in susceptible bacteria given a functional form *F*(*P)* that varies with phage. For T7, *F*(*P*) = *ϕP* with adsorption rate *ϕ* while for T4, we use a nonlinear profile 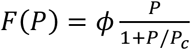 with a saturating phage density *P*_c_. Phage saturation mimics Michaelis-Menten reaction kinetics effectively mediating inter-phage competition processes such as lysis inhibition^46^, prevalent in T-even phage^49^. Infected cells progress through *L* infection stages *E*_*i*∈{1…*L*}_ before lysis – these stages are a mathematical convention to represent latent periods as an Erlang distribution^41^ with an average latent period 1/*η*(*R*) and Coefficient of Variation 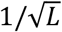, generalizing the *L* = 1 scenario corresponding to exponentially distributed latent periods. Lysis produces *β*(*R*) virions. We assume that the lysis rate 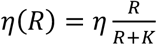 and burst size 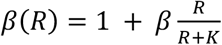 depend on growth rate such that phage lysis halts in the absence of available resources and lysis (when it occurs) has a minimum burst size of 1. Finally, phage decay at a rate *ω* per virion. All phage reaction terms, are modulated by a nonlinear function 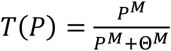, where 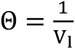 is a discreteness threshold scaling as the inverse of the volume of the PDE integration lattice sites *V*_*l*_ (see Supplementary Text for numerical implementation), with large Hill coefficient *M*. This nonlinear term sharply drives phage rates to 0 when phage densities are lower than 1 virion per lattice site *P* < Θ, and ensures that phage, the fastest dynamical variable in the system, do not diffuse and subsequently infect cells before they reach biologically realistic densities (see Supplementary Text).

<u>Advection terms – chemotaxis:</u> Bacteria sense attractant gradients and move chemotactically towards higher concentrations given the self-generated velocity field 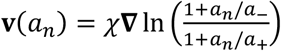, which triggers a collective population range expansion wave^36,37^. The sensing function takes a logarithmic Weber form^36,50^ with two sensitivity parameters *a*_−_ and *a*_+_ respectively denoting the lowest and the highest chemoattractant densities bacteria react to. The scalar χ, modulating the velocity magnitude, represents bacteria responsiveness to sensed chemical gradients. For simplicity, we assume that the velocity field function does not depend on the attractant type *n* through the parameters *a*_−_, *a*_+_ and χ. Infected cells also sense differences and move chemotactically towards higher concentrations with a potentially different efficiency than uninfected cells.

### Chemoattractant-mediated infection-chemotaxis feedback drives dynamical stability of PhASERs

We leverage PDE model simulations to explore feedback mechanisms responsible for PhASER emergence, persistence, and stability. Figure 3A shows timepoints from a simulation where bacteria, sensing a single attractant, give rise to a single range expansion front^36^, with zoomed-in insets in the second row. When the front encounters and is invaded by phage, a lysis region spreads out before the fronts regions on either side slowly curl back inwards and converge, just like in the experiments shown in Figure 1. The first inset in Figure 3A (first panel in the second row) shows that this convergence is driven by an infection-induced curvature in the attractant gradient (green isolines) that arises from reduced consumption due to phage killing of host cells in the center of the front. The bacteria migrate orthogonally across the isolines in the direction of the chemotactic flux (see arrows in Figure 3A). At the same time, infected cells (blue colormap), which migrate together with non-infected ones, lyse and release new free phage (red colormap in Figure 3A) close to the front. Susceptible cells moving inward interact with free phage and transport them during infection as shown by the change in color of the flux arrows, denoting the fraction of infected cells. Once the two wings of infection transport reunite, they influence one another’s chemotactic collective behavior through the chemoattractant profile, which merges the attractant isoline, forming a cusp (second inset in Figure 3A, second row). As a result, the cells at the merging point exhibit a high infection prevalence and accumulate along the symmetry line. The lagging, cusp-shaped front produces a steeper chemical gradient and therefore a faster migration in the center of the front-associated infection zone (Figure 3A, third inset). This population-driven chemical feedback allows infected cells to migrate forward faster ensuring the dynamical stability of PhASER propagation.

**Figure 3.**
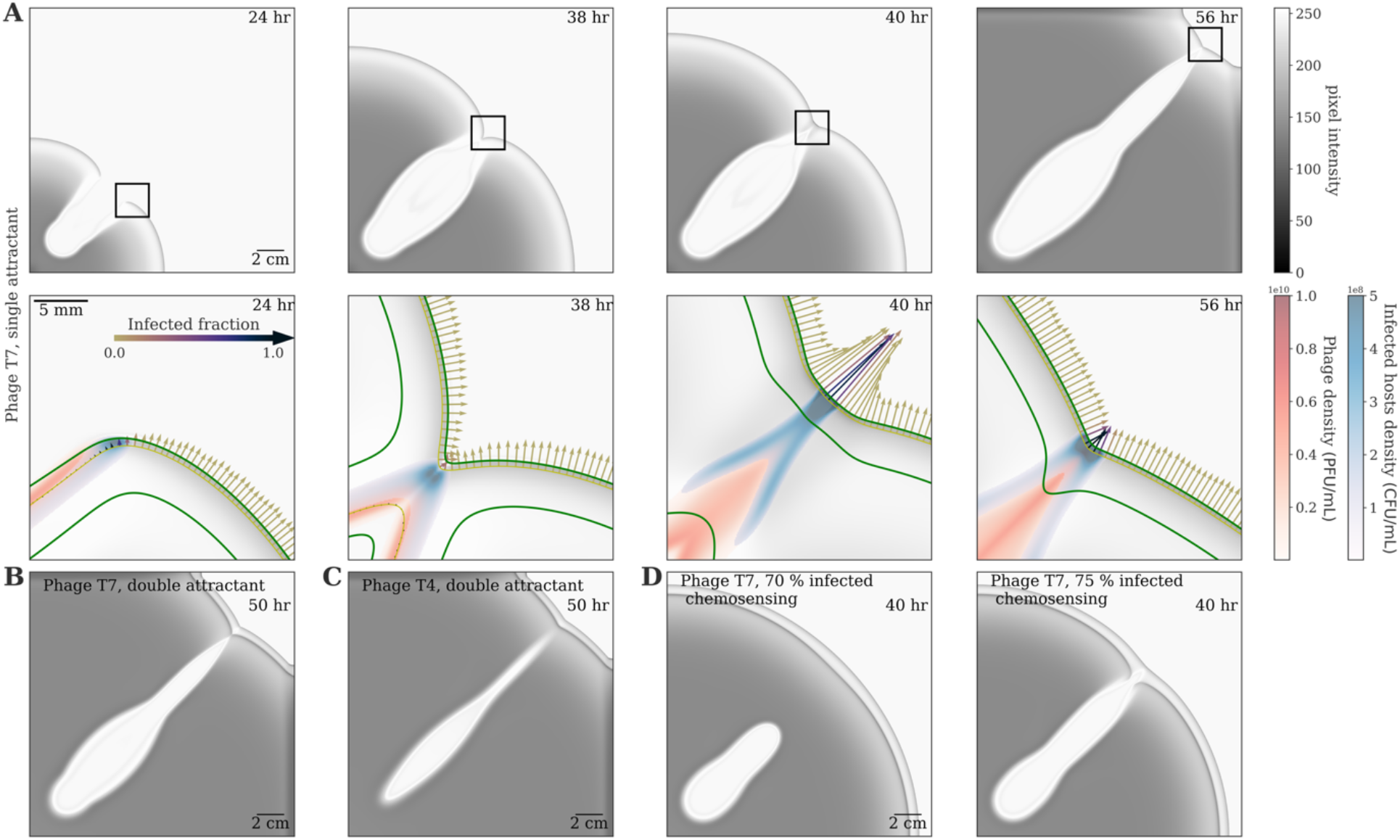
Model of phage infections transport by chemotactically migrating bacteria populations. **(A)** Timepoints of a simulation using the model defined in Box 1, with parameters inferred for *E. coli* W1485(F8) and phage T7 (see Supplementary Text) and a single attractant (yielding a single chemotactically migrating front). Pixel intensity is mapped from calibrated cell density (Methods). The macroscale spatiotemporal dynamics (top) closely resemble the patterns of Fig.1A. Bottom: insets showing density of total (grey) and infected (blue) cells and density of phages (red). The two green lines represent attractant density isolines at half the initial value and at the lower sensing threshold. The thin yellow line, close to the front, denotes the isoline where the chemosensing response is at 80% its maximum. Quivers represent the chemotactic flux along this line, colored according to the fraction of infected cells. **(B)** Snapshot of the same simulation as **A**, with the addition of a second attractant. The qualitative behavior (invasion transient and coalescence to a stably propagating infection) of the model is not affected by the resulting dual migration fronts. **(C)** Simulations with viral life-traits inferred for phage T4, while maintaining all bacterial traits unaltered (see Supplementary Text). The model behaviour transitions to a macroscale pattern closer to the empirical T4 PhASER shown in Figure 1D. **(D)** Accounting for phage-inhibited chemotaxis of infected cells. Model simulations are shown for W1485(F8) encountering phage T7, while modulating infected cell responsiveness to attractant concentrations. The stable infection transport phenomenon persists for a 25% reduction in chemotactic responsiveness of infected cells, but is finally lost as responsiveness is lowered to 70% that of uninfected cells.

After some time, the dynamics stabilize to a steady state where the infection transport remains unaltered as the wave progresses, as indicated by the conserved flux profiles across the front in the fourth inset of Figure 3A. The infection is focused by the convergent migration of cells up the nutrient gradient toward the symmetry line. Infected cells then migrate with the expansion front, eventually lysing and producing free phage which in turn infect new cells in the front ensuring a long-range stable infection transport. The control of bacterial densities due to lysis maintains slower chemoattractant consumption in the infected front region. This slower consumption changes the curvature of the gradient such that the fronts of uninfected cells bend inwards, ensuring a steady influx of susceptible cells into the front region where phage infection is active. This convergent migration towards the symmetry line also focuses the infection and constrains its lateral spread. We stress that such range expansion is a population-driven collective behavior akin to chemotaxis of bacteria in the absence of phage^36^. Therefore should a few cells (infected or not) move ahead of the front due to random fluctuations in their chemotactic process these cells will not generate a sufficient attractant gradient to sustain range expansion and the fluctuation will dissipate^51^.

The processes stabilizing PhASERs with a single chemoattractant molecule also apply to simulations including a second chemoattractant molecule, which produces a second chemotactically migrating front (Figure 3C), similar to that observed for W1485(F8) and its W1485 ancestor in LB swim agar (Figure 1D & Supplementary Figure 2). Finally, we modified the phage life-history traits in model simulations to correspond to phage T4 (including the infection rate, maximum lysis rate, and burst size) while keeping the bacteria parameters fixed. Model simulations can qualitatively recapitulate experimentally observed T4 dynamics (compare experimental observations in Figure 1D with simulations in Figure 3C), generalizing our finding for model simulations with a single front (as shown in Figure 2).

### PhASER initiation is robust to infection-reduced chemotaxis

On infecting a host cell, lytic phage can redirect cellular machinery, resources, and energy stores to the rapid production of phage progeny^52–54^. The viral-induced reshaping of bacterial metabolism and physiology could also impact bacterial chemotaxis – which depends on active gene expression, continuous energy expenditures and cell structural integrity^46,55^. Therefore we leveraged our model to evaluate whether phage-induced attenuation of chemotaxis efficiency could lead to infected cells falling behind the front and, in turn, PhASER termination. We hypothesized that there would be a critical threshold below which reduction in the chemotactic efficiency of infected cells would lead to PhASER termination. We tested this hypothesis in the reaction-diffusion-advection model by assuming that infected cells perform chemotaxis with responsiveness 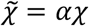 with *α* between 0 and 1 relative to uninfected cells. Note that in our formalism this parametrization is equivalent to a scenario where, at any point in time, a fraction *α* of the infected cells move chemotactically. As expected, when simulated phage infections severely limited cells’ chemotactic efficiencies, infections fell progressively behind and were lost at the back of the range expansion front. Surprisingly, we found that the system was remarkably tolerant to reduction of infected cell chemotaxis. Maintenance and infection transport of a T7-parameterized PhASER occurred above a chemotactic responsiveness threshold between 70% and 75% that of uninfected cells (Figure 3D). In comparison, in the absence of phage, similar chemotaxis deficits in bacterial subpopulations produce significant disadvantages, as less chemotactically responsive bacterial mutants rapidly lose their position in migrating chemotactic fronts^56^. Note that the threshold for infection transport depends on the phage infection parameters. For example, phage T4-parameterized PhASERs maintain the ability to propagate only if chemosensing responsiveness exceeds 85% to 87% relative to uninfected cells (see Supplementary Figure 12). This finding reinforces the central role of infected cell migration in the generation, stability, and persistence of PhASERs.

### PhASERs are robust to large-scale environmental perturbations

We set out to assess the robustness of PhASERs to large-scale environmental perturbations. First we modulated the agar density, which impacts the movement of cells in swim plates, slowing the progress of chemotaxing bacteria with increasing agar concentration. We found that stable infection transport was robust to a wide range of agar concentrations, corresponding to dramatic differences in range expansion speeds from 3.5 to 8.5 mm/hr (Fig 4A). The model reproduces this robustness, as we could adapt the chemosensing responsiveness *χ* and the bacteria diffusion constant *D*_*B*_ to reproduce the PhASERs geometries at the experimentally measured range expansion speeds. We note that when the agar concentration is sufficiently high it can disrupt regular chemotactic range expansion which in turn leads to the failure of stable infection propagation (Supplementary Figure 13). Moreover, the numerical exploration in Supplementary Figure 14 suggests that our PDE model’s qualitative dynamics are robust to perturbations in nutrient quality. As one example, given enhanced nutrient availability, PhASER propagate in a narrower lysis region at range expansion speeds of ∼15 mm/hr – comparable to chemotactic front speeds reported in liquid media^57^.

**Figure 4.**
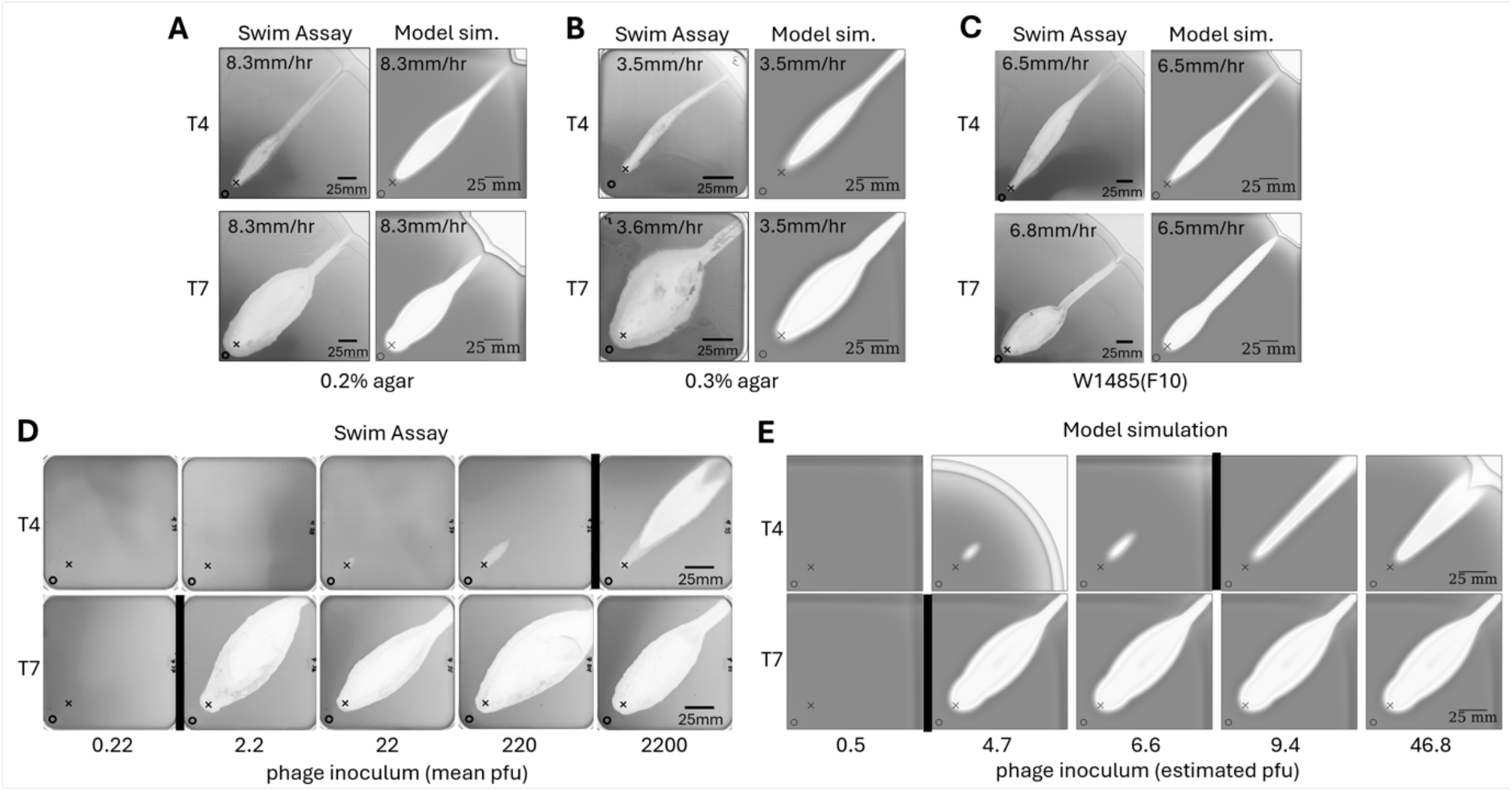
Stable phage infection transport by chemotaxing bacterial populations is robust to environmental and host changes that impact migration speed, and in phage-dependent manner, to invading phage population size. **(A)** Reducing agar concentration in swim assays from 0.25% (Fig 1A) to 0.2% increases *E. coli* W1485(F8) migration rate by ∼50%. However, invasions of hosts migrating from location ○ by phages T4 (top) and T7 (bottom) inoculated at location × on 24x24cm assays continue to resolve to stably propagating infections, Experimental data (blue channel of transmission scans) to the left are closely mirrored on the right by model simulations (cell concentration-calibrated blue pixel values) in which host parameters *χ* and *D*_7_are adjusted to match the front expansion speed (see Supplementary Text Section 2) while T7 and T4 phage parameters are unchanged, showing that the model ingredients encode the observed empirical robustness. **(B)** Similarly, increasing agar concentration to 0.3% slows host migration by approximately 50%. Yet, phages T4 and T7 invade and produce stable PhASERs on 12x12cm plates, indicating that localized phage invasion and stable infection transport in migrating bacterial fronts is robust across a range of environmental perturbations to migration speed. **(C)** T4 and T7 PhASERs robustly emerge on *E. coli* W1485(F10), a spontaneously evolved fast migration mutant of W1485(F8) that maintains two chemotactic fronts (left). Model simulations (right) where the bacterial parameters are quantitatively fit to the new strain (see Supplementary Text Table S1) reproduce the experimental behaviour. **(D)** Effect of the initial phage concentration on PhASER formation is measured by challenging *E. coli* W1485(F8) migrating from location ○ with few to thousands of plaque forming units (pfu). T4 requires needs ∼1000 phage particles to invade and initiate infection transport, while just a few phage T7 particles reliably initiate stable PhASERs (cutoffs indicated by black verticals). **(E)** The qualitative scenario of phage infective dose dependence is reproduced by a modified model, although the quantitative transition from T4 loss to stable infection propagation is steeper than observed in experiment. Critically, the phage inoculum dependence was not reproduced by our original model. Reproducing the qualitative experimental observation required a reduced chemosensing responsiveness of 87% for infected cells, reflecting partial suppression of host chemotactic motility by phage infection. The PFU count, expressed in units of the discreteness density threshold Θ, is estimated right before the range expansion front encounters the phage inoculum. Each column, from left to right, corresponds to an initial phage density of 5 · 10^4^, 5 · 10^5^, 7 · 10^5^, 10^6^ and 5 · 10^6^ PFU/mL.

Second, we explored the impact of mutant propagation speed on PhASERs. We do so because in migrating bacterial systems there is a particularly strong selection for mutants with enhanced front propagation speeds^56^. Here, we asked whether naturally occurring migration speed mutants of our ancestor bacterial strain would potentially disrupt PhASER formation and stability. We observe that as in the case of the faster expanding W1485(F9), a mutant W1485(F10) that gives rise to much faster range expansion than its parent W1485(F8) but with two chemotactic fronts, nonetheless reliably produces PhASERs both in simulations and in experiments (Fig 4B). This finding agrees with the observation that mutations affecting the number of range expansion fronts (i.e., single or double) does not noticeably impact PhASER emergence, propagation, and stability (Fig. 2).

Finally, we examined how the size of the phage inoculum impacted its invasion of the migrating bacterial front and the emergence of stable, long-distance infection transport. To do so, we challenged migrating bacterial fronts with phage inoculi containing either phage T4 or T7 at numbers of plaque forming units (PFUs) spanning several orders of magnitude. Strikingly, we observed that just a few PFUs of T7 reliably invade the host front and initiate PhASERs (see experiments and model in Figure 4D,E respectively). In contrast, phage T4 produced increasingly extended plaques with larger inoculi but was unsuccessful in invading the migrating front and initiating stable infection transport below a threshold level between 10^3^ and 10^4^ PFUs. We posit that this Allee effect^58^, mediated by host population range expansion, is driven by a requirement for a sufficient initial killing-induced deformation of the chemotactic front, enabling a sustained influx of susceptible cells into an infection region and establishment of a proliferating phage population, leading to the stable PhASER. Our model can reproduce this qualitative trend showing the absence of an Allee effect for T7 parameters and the presence of an Allee effect for T4 parameters, (albeit, at a different inoculum range than experiments), when T4 infected cells have slightly reduced chemotactic efficiency (Fig 4E, Supplementary Figure 15). By comparison, such Allee effect is absent from simulations when the chemotaxis of T4 infected cells is at full efficiency (Supplementary Figure 15 A), or when the chemotaxis of T7 infected cells is reduced by a similar amount. This result indicates that this qualitative model trend is driven by a combination of chemotaxis reduction and phage life-history traits, although we cannot exclude the possibility of Allee effects independent of infection-reduced chemotaxis for every parameter combination associated with distinct phage-host pairs.

### Dynamical robustness of interactions between PhASERs and with other migrating populations

In order to probe the infection transport stability of PhASERs to dynamic perturbations we set out to investigate collisions of PhASERs with migrating bacterial waves. We first grew two colonies from different locations, each transporting a PhASER, towards a common location. When equivalently translocating PhASERs collide at a shallow (60 degree) angle, they merge in their average direction (Fig. 5A). Numerical simulations of our model reproduce this phenomenon without additional parameter changes and reveal that this dynamical stability is enabled by precisely the same feedback that permits stable coalescence of the two sides of the invasion transient that produces T7 PhASERs. Simulations reveal that as the two outer fronts of the PhASERs merge, the infected cells along the symmetry line are propelled faster into the uncolonized region by the attractant gradient cusp and by the presence of more cells at the colonies merging interface (Fig. 5B, Supplementary Movie 6). In contrast, when we repeated the experiments with two PhASERs merging at a much steeper (120 degree) angle, the collision resulted in a termination of the transport process both in the experiments and in the model (Fig. 5C). When the relative angle of two approaching fronts is too steep, the radial portion carrying phage is lost in the bulk behind the resulting front, as the phage-free sectors of the expanding populations merge and close the nutrient-rich corridor faster than the infections can travel to keep up with the front (Fig. 5D, Supplementary Movie 7). Similar outcomes are obtained when PhASERs collide with walls in the environment at different angles (Supplementary Figure 16). Together, these results reinforce the importance of geometry underlying robust ecological dynamics^59^. Finally, we tested the stability of infection transport to complex ecological interactions featuring competition between different host traits. We merged a colony of *E. coli* strain W1485(F8) carrying a T4 phage PhASER with a significantly faster colony of *E. coli* strain W1485(F10) (experimental front speeds of 4.5 mm/hr vs. 6.5 mm/hr, respectively). Despite the mismatched hosts, the propagating phage infection transfers to the faster strain, continuing to stably propagate on the curved interface where the two populations merge (Fig 5E, Supplementary Movie 8). This behaviour is closely reproduced by the model without needing to adjust any parameters, underscoring the generality of predictions within our theoretical framework (Fig 5E, Supplementary Movie 9), including robustness to differences in phenomenological representations of phage adsorption (Supplementary Figure 17).

**Figure 5.**
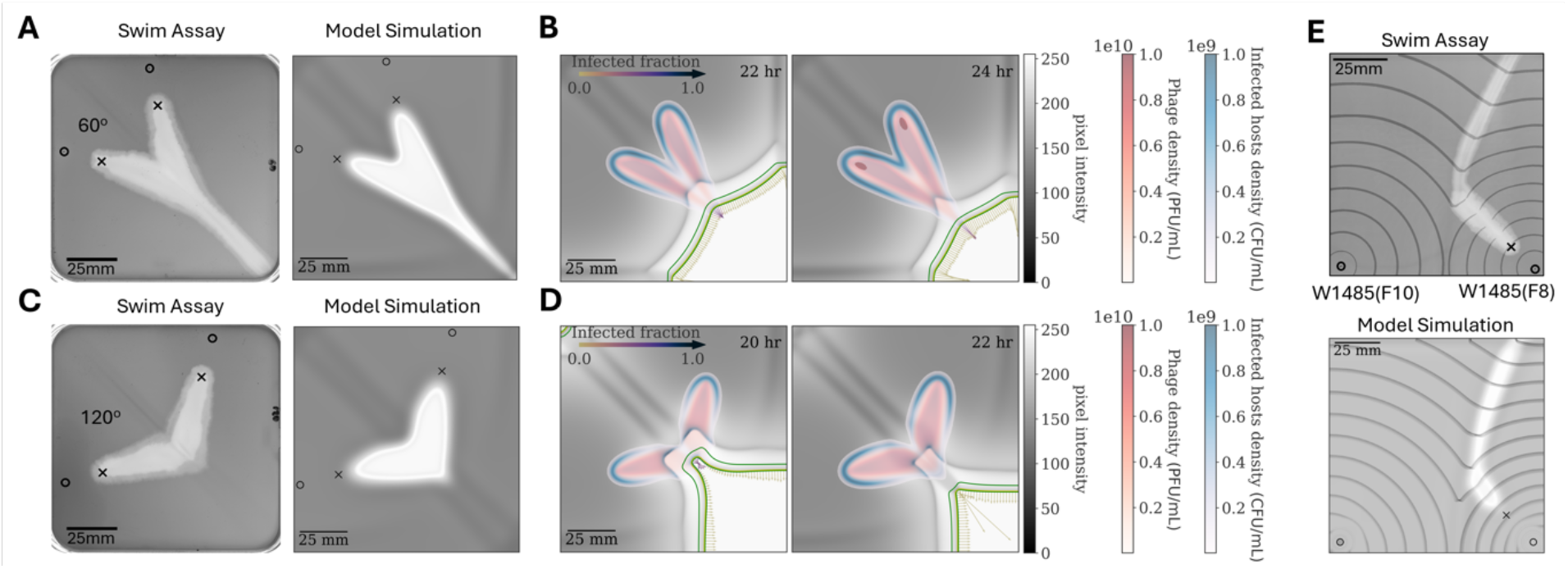
Geometric dependence of merging and termination of PhASERs on colliding chemotactic waves. **(A)** Experimental image (left) shows two T4 PhASERs on nearby *E. coli* W1485(F8) colonies that migrated from location ○ and intersected at a 60 degrees angle. The PhASERs merged along the symmetry line (the plate’s diagonal) and continued stable progression across the agar plate. This phenomenon is reproduced by the model (right). The parameters correspond to those inferred for *E. coli* W1485(F8) and phage T4. **(B)** Detailed simulation output (see Fig. 2A bottom) at two timepoints shows PhASER merging is driven by increased flux of infected cells along the symmetry line, mediated by attractant profiles, similar to phage T7 stabilization in Figure 2A (Supplementary Movie 6). **(C)** Increasing the intersection angle to 120 degrees halts PhASER infection transport the two colonies collide, both experiments (left) and simulations (right). **(D)** Detailed simulation timepoints show that when the angle between the fronts is too steep, the two colliding fronts merge at a point beyond the phage infections, which are lost behind the front as attractant and nutrients are depleted (Supplementary Movie 7). **(E)** Experiments (top) and simulations (bottom) where a T4 PhASER on *E. coli* W1485(F8) collides with migrating front of a much faster *E. coli* strain W1485(F10), turns sharply and continues in the faster curving interface between the strains (See also Supplementary Movies 8,9). The PhASER transport phenomenon is sufficiently robust that the infection transport dynamically adapts to higher speeds, for instance encountered upon the emergence of a faster mutant, and this robustness is reflected in the model. The simulation parameters are those inferred for phage T4 against W1485(F8) and W1485(F10) respectively (see Supplementary Text Table S1). Time frames are overlaid with 2 hour-spaced front positions (grey curves).

## Discussion

Long-range, stable and robust transport of infections spontaneously emerges from interactions between virulent phage and migrating bacterial hosts, including in multiple strains of *E. coli* and *P. aeruginosa*. We find that phage interacting with a migrating chemotactic front initiate an early-stage lysis expansion and then transition to a stable linear infection transport regime (*i*.*e*., ‘PhASERs’). These PhASERs propagate over scales well beyond observations of localized transport phenomena in prior work^38–40^. To explore the basis for PhASERs, we developed and analyzed a PDE model which builds upon prior studies of both chemotactic and virus-microbe dynamics^17,36,39^. This model successfully reproduces the emergence, stability, and robustness of PhASERs. Together, our joint theoretical and empirical investigation shows how local phage killing reduces nutrient consumption, establishing a concave nutrient gradient that draws in susceptible hosts and projects infected cells to an infection tip in a migrating bacterial front. Our model revealed that this population-induced chemical feedback has a crucial role not only in the emergence and long-range persistence of phage infection transport, but also on its stabilization upon dynamical perturbations. Moreover, our experimental and simulation findings jointly suggest that the emergence and stability of phage infection transport over long distances is enabled by chemotaxis of infected cells. We do note that PhASER dynamics are not inevitable. Our results reveal that phage-host traits and environmental factors can shift outcomes to bacterial chemotactic escape or local host extinction.

The present study of stable and robust long-distance phage infection transport with chemotactic bacteria populations comes with caveats. Here we focused on the leading edge of phage and bacterial transport while neglecting coevolutionary dynamics of phage resistance by bacteria and counterdefense mutations by phage. In prior work over short distance scales, phage-bacteria co-propagation was found to be resilient to coevolution^40^, yet rare mutations or specific initial spatial configurations could destabilize infection transport^39^. We anticipate that the inclusion of coevolutionary dynamics will be particularly relevant behind the migrating front where phage-resistant bacteria would have a significant advantage and counter-defense mutant phage could potentially recolonize. Assessing the impacts of eco-evolutionary dynamics in both theory and experiments offers a promising direction for future studies. Doing so would also help to shed light on mechanisms of phage and bacterial coexistence that have been perceived to be fragile given asymmetries in coevolutionary-induced adaptations that favor bacterial escape from infection^10^. Here, the presence of an advancing front acts to maintain a source pool of host bacteria for virulent phage, even as the majority of the bacterial population remains out of reach. Broadening the ecological relevance of the work will also require considering additional model systems and infection modalities, including the potential transport of temperate infections where viral genomes integrate into the hosts.

Investigating long-range infection transport within (controlled) *in-situ* environments will also help elucidate the impact of spatial heterogeneities and confined geometries on phage-bacteria coexistence. We also stress that our theoretical analysis leverages the large populations involved in range expansion, where demographic stochasticity minimally affects growth-death dynamics. By contrast, the outcome of lytic phage infections are variable^44,60^. Stochastic resource-mediated infections can generate heterogeneity in phage transport, particularly from specific spatial virus-host configurations starting at low densities^61^. In the future, developing stochastic models will be essential to quantifying this variability and its consequences for infection propagation. At the same time, new experimental approaches are needed to quantify how specific infections alter cell functions^53,54,62^, e.g., by tracking the chemotactic behavior of infected hosts, whose motility shifts may influence population-scale dynamics driving PhASER emergence and stability. Linking cellular outcomes to population-scale dynamics will also help clarify the extent to which phage can co-propagate with expanding bacterial populations in the context of novel colonization dynamics across different environments.

In summary, combining theory and experiments, we characterized the emergence of a robust balance between infection propagation and chemotactic escape in spatial environments, enabling long range transport of bacteriophages and stable coexistence in the absence of coevolutionary arms races. We envision that this study will help pave a path towards integrating eco-physical and eco-evolutionary mechanisms underlying phage-bacteria coexistence in complex habitats.

## Methods

### Strains

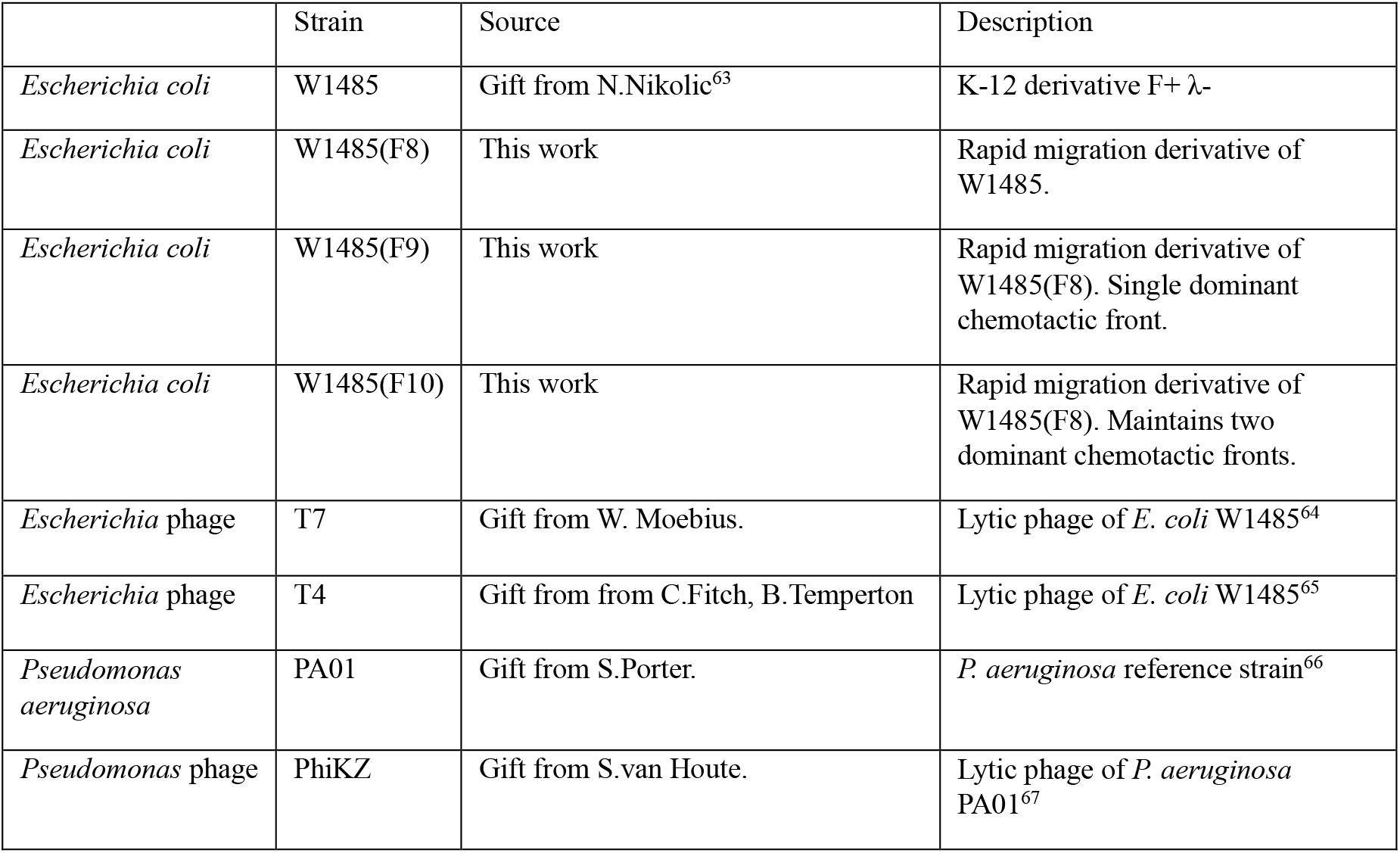

|  | Strain | Source | Description |
| --- | --- | --- | --- |
| *Escherichia coli* | W1485 | Gift from N.Nikolic<sup>63</sup> | K-12 derivative F+ λ- |
| *Escherichia coli* | W1485(F8) | This work | Rapid migration derivative of W1485. |
| *Escherichia coli* | W1485(F9) | This work | Rapid migration derivative of W1485(F8). Single dominant chemotactic front. |
| *Escherichia coli* | W1485(F10) | This work | Rapid migration derivative of W1485(F8). Maintains two dominant chemotactic fronts. |
| *Escherichia* phage | T7 | Gift from W. Moebius. | Lytic phage of *E. coli* W1485<sup>64</sup> |
| *Escherichia* phage | T4 | Gift from from C.Fitch, B.Temperton | Lytic phage of *E. coli* W1485<sup>65</sup> |
| *Pseudomonas aeruginosa* | PA01 | Gift from S.Porter. | *P. aeruginosa* reference strain<sup>66</sup> |
| *Pseudomonas* phage | PhiKZ | Gift from S.van Houte. | Lytic phage of *P. aeruginosa* PA01<sup>67</sup> |

### Bacterial and phage stocks

All bacterial strains were grown overnight from single colonies to saturation, mixed with glycerol to a final concentration of 17% and stored at as a master stock and single-use aliquots (at concentrations of approximately 10^9^ cfu/ml) -80ºC. Single-use aliquots were thawed for inoculation of liquid cultures or direct inoculation of soft agar swim assays. Viable cell concentrations in stocks were determined by colony counts of thawed, serially-diluted samples.

Phage were amplified by adding cores picked from single plaques on double layer plates to growing host cultures (at 10^7^-10^8^ cfu/ml). After several hours, clear lysates were centrifuged to remove cell debris and the supernatant was passed through a 0.2µm syringe filter and stored in amber glass vials at 4C. *E. coli* phages were amplified on *E. coli* strain W1485. Phage PhiKZ was grown on *P. aeruginosa* PA01. Plaque forming unit concentrations were determined by serial dilution and plaque counts on double overlay plates^68^.

### Host migration training for *E*. *coli*

Soft agar swim assays select for faster migration at the chemotactic front. As faster migrating mutants of *E. coli* W1485 emerge regularly enough to disrupt our swim assays, we passaged this strain across a series of swim assays to obtain more stable, faster migrating derivatives for our experiments^69^. For each passage, we inoculated the most recent isolate at a corner of the plate and transferred the first arrivals at the opposite corner to the next plate. The faster, more stably migrating W1485(F8) was obtained after eight passages. W1485(F9) with a single dominant chemotactic front, and W1485(F10), retaining two fronts, emerged on a ninth passage, and migrated at a similar rate to each other. *P. aeruginosa* PA01 did not show rapid accumulation of fast migration mutants and was not passaged.

### Scanner timelapse platform

Two scanner platforms, based on a previously reported design^70^ were used to record time-lapse image sequences of swim assays. Each platform includes three Epson V800/850 scanners, that can collect large format (8”x10”) transmission scans. The scanners are connected to a Raspberry Pi 4 computer (Supplementary Figure 18) and controlled by a custom python script (Supplementary Method 1) that uses Scanner Access Now Easy (http://www.sane-project.org/) and a Sane python interface (https://pypi.org/project/python-sane/) to collect images from all running scanners at user-determined intervals and resolutions. Custom inserts (Supplementary Method 2) are used to align assay plates in the transmission imaging region of the scanner platens.

### Swim assays

All swim assays were performed in 12cm (Greiner, 688161) or 24cm (Corning CLS431301) square petri dishes. Low concentration swim agar media was prepared by combining warmed LB (10 g/L tryptone, 5g/L yeast extract, 5g/L NaCl) liquid broth with premelted LB containing 2% Bacto Agar (Difco) in appropriate proportions at 50ºC. Volumes (50ml or 200ml) of mixed media were added to 12cm or 24cm plates, respectively, and allowed to set. Host and phage stocks were inoculated by pipetting (typically) 1µl of each directly into the agar at predefined locations indicated on a map beneath the plate. Plates were then left for 1 hour to absorb the inocula and overlaid with clear mineral oil (Merck, 330779) to prevent evaporation and condensation on lids. Once overlaid, the plate lids were dried of condensation and plates incubated (positioned on transmission scanners if timelapses were to be recorded). Typical timelapses recorded 300dpi colour images at 10 minute intervals.

### Image pixel intensity to CFU/ml calibration

We obtained an empirical calibration curve to allow inference of cell concentrations in the swim agar from pixel intensities in our scanned images. To do so, we performed 2-fold serial-dilutions from an overnight (37ºC, 230RPM) culture of *E. coli* W1485. We added 50ml of each dilution to 12x12cm swim assay plates (to match light path length in swim assays). The plates were immediately scanned and fit obtained for media blank-subtracted average pixel levels (blue channel) as a function of corresponding cell concentrations determined by colony forming unit counts. See (Supplementary Figure 19).

### Image analysis

Primary image analysis of timelapse images was performed using FIJI software^71^. The blue channel of the colour scan was analyzed as it showed a slightly higher dynamic range in detecting cell density than the red and green channels. Time differential image sequences, extracted from the timelapse image stacks by subtracting each image from the one preceding it, were used to identify cell growth regions and particularly to highlight rapid density changes at migration and lysis fronts (Supplementary Movie 10). Time differentials were binarized to extract migration fronts. Binarized images were denoised with median and morphological close/open filters. Maximum projection of binarized front images through time was used to generate overlays of time-spaced front shapes throughout PhASER swim assays. (Supplementary Figure 3). Front migration rates are determined by displacement of advancing chemotactic fronts in kymographs (obtained using the FIJI multi-kymograph plugin) and conversion to mm/hr units (Supplementary Figure 20).

## Data Availability

The datasets generated and/or analysed during the current study are available as image and movie files shown in the Main Text and in the Supplementary Information. All custom code used in acquiring and analyzing the data is available to reviewers and will be made available upon publication via github and zenodo.

## Acknowledgements

We are grateful to B. Temperton, T. Bergmiller, N. Nikolic, W. Moebius, E.S. Tamar, R. Kishony, I. Golding, S. Pagliara, A. Andersson, N. Stitt, M. Cooper, I. Parry, and the members of Exeter’s Bacterial Pathogenicity Research Group and University of Maryland’s Quantitative Viral Dynamics Group for helpful discussions and suggestions. We thank N. Nikolic, C. Fitch, B. Temperton, S. Porter, S. van Houte, R. Sorek, W. Moebius for kind gifts of strains and Exeter’s Physics Machine Shop and Microfluidics Facility for technical assistance. The research leading to these results was made possible via funding from Exeter Biosciences (RC), EPSRC Summer Internship (CK), Exeter Access2Internship (CK), Simons Foundation 930382 (JSW) and support from the Chair Blaise Pascal program of the Île-de-France region (JSW, JM, and AG).

---

**Box 1 Figure.**
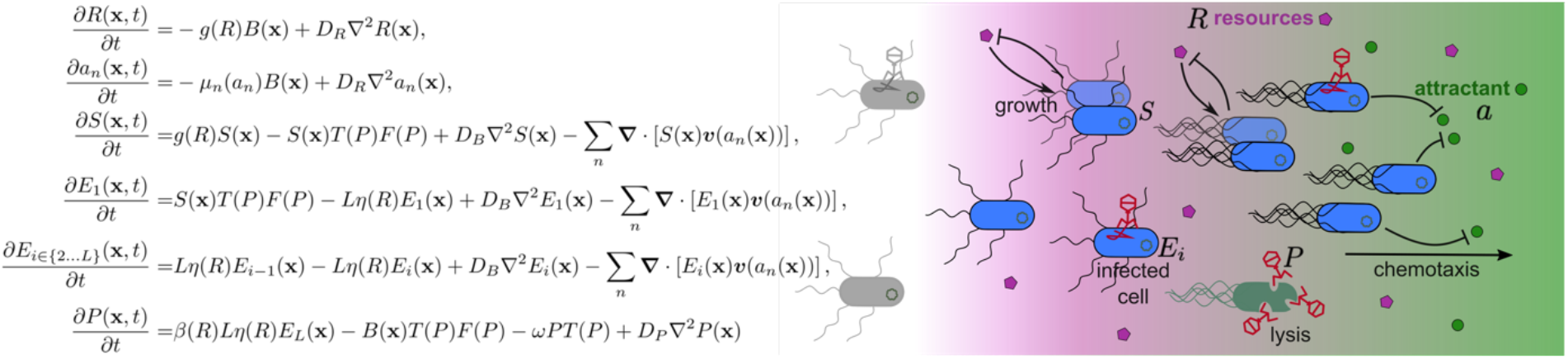
**Caption:** Spatial PDE model governing the spatiotemporal concentrations of chemotaxing susceptible (*S*) and phage-infected (*E*_i_) bacteria, phage *P*, resources *R* (magenta gradient) and a set of chemoattractants a_n_ (green gradient). Bacteria feed on resources at a rate *g*(*R*) and consume attractants at a rate µ_*n*_(*a*_*n*_), generating a spatial chemical gradient. As a result, bacteria move chemotactically towards higher attractant concentrations through the self-generated velocity field **v**(*a*_*n*_), triggering a collective range expansion wave. Free phage infect bacteria at a nonlinear rate *F*(*P*), modulated by a Hill function*T*(*P*) incorporating virion discreteness effects. Infected cells continue to move chemotactically while they complete *L* infection stages over an average latent period 1/η(*R*), after which cells lyse producing β(*R*) virions. Both η(*R*) and β(*R*) decrease as a function of resources *R*, slowing down phage infections. Finally, phage are depleted at a rate ω*T*(*P*). All agents diffuse according to a specific diffusion constant *D*.

