## Supplementary Figures for "Stable coexistence and transport of lytic phage infections with migrating bacterial hosts"

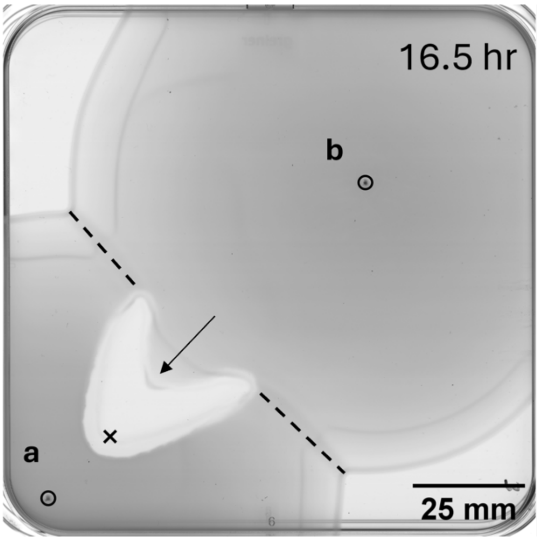


**Supplementary Figure 1.** A phage-protected but infection-free region can exist within a phage invasion zone. Two *E. coli* W1485(F8) colonies (a,b), migrate outwards through LB swim agar from locations marked by ∘. Colony a, encountering T7 phage at location × develops a clear phage protected zone in the advancing front. As migration halts on collision of colonies a and b (dashed line), the front advancing from b penetrates between the infected wings and into (arrow) the phage invasion zone without lysis before being halted at the periphery, illustrating that phage infection and production can be restricted to the invasion zone edges.

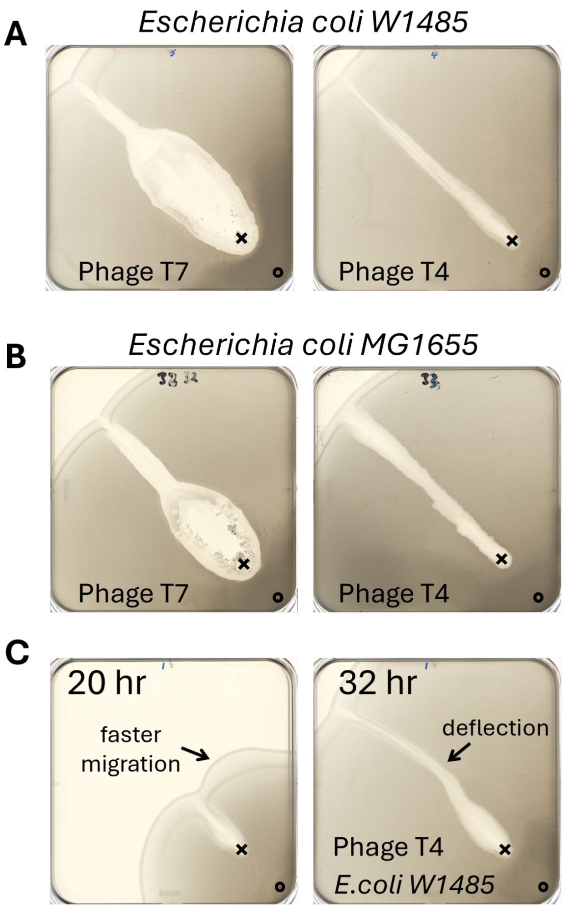


**Supplementary Figure 2.** Phage T7 and T4 PhASERs on *E. coli* strains W1485 and MG1655**.** *Escherichia coli* strains **(A)** W1485 and **(B)** MG1655 migrating through 12cm square LB agar swim plates both form PhASERs on encountering phages T7 (left) and T4 (right). Host cells and phages are inoculated at locations **o** and **x**, respectively. **(C)** Local emergence of faster migrating cells appears first as a spreading bulge in an *E. coli* W1485 front at 20hr (left), which deflects the T4 PhASER (right). Agar concentrations are 0.25% in **A,C** and 0.3% in **B**. Swim assays are performed at 30C as otherwise described in Methods.

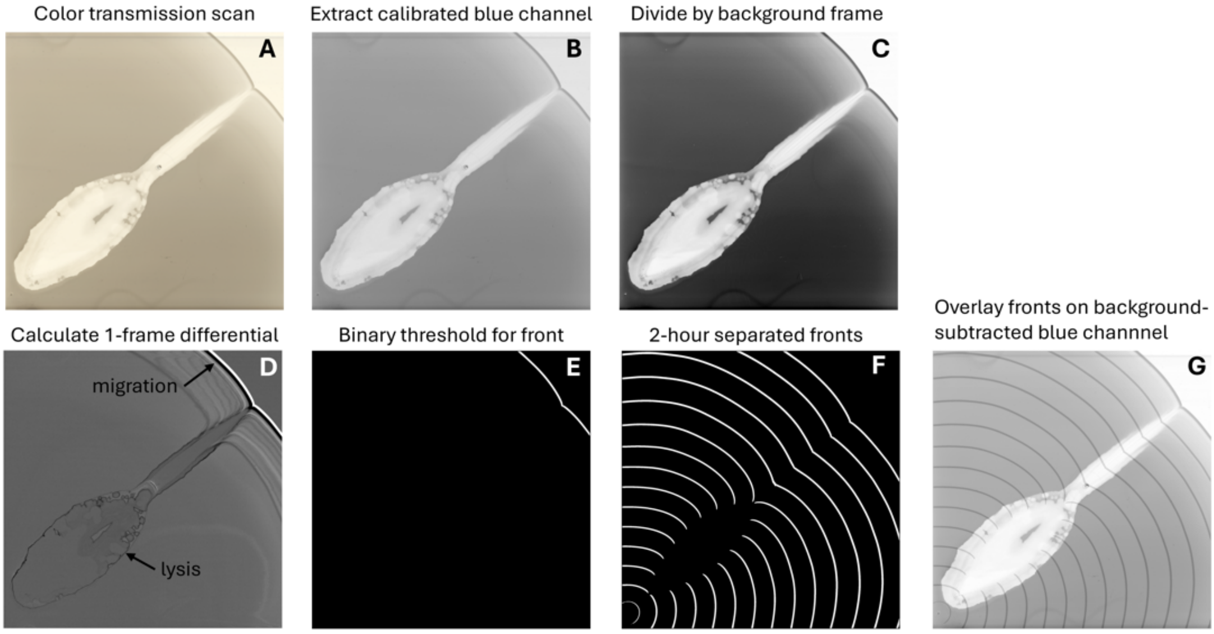


**Supplementary Figure 3.** Sample image processing steps for timelapse scans of migrating bacteria intersecting phage inocula. Images are processed to highlight spatial cell density and change due to cell growth, migration and lysis. For *E. coli* W1485((F9) interacting with phage T7 (as Figure 2), a scanned transmitted image sequence **(A)** is separated into colour channels. The blue channel **(B)** is calibrated to cell concentration (Supplementary Figure 9). The blue channel is normalized **(C)** by division by the first (background) frame to reduce lighting inhomogeneities. Single frame differential images **(D)** are obtained by subtracting consecutive frames and reveal adjacent light/dark bands of migrating fronts, light background cell growth, and dark edges indicating cell killing. Binary thresholding **(E)** captures the migrating bacterial front. Maximum projection of 2 hour separated fronts **(F)** reveals the position and trajectory of the interaction across time, and can be overlaid on cell image (**G**, Figure 2). Elements of this analysis (**A,C,D,E**) throughout a timelapse sequence are shown in Supplementary Movie 10.

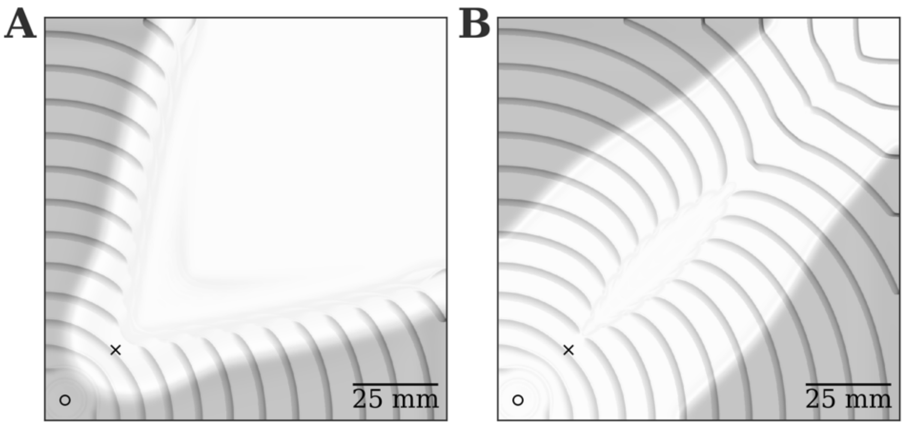


**Supplementary Figure 4.** Simulated dynamics of a T7 PhASER carried by *E. coli* W1485(F8) bacteria (host and phage inoculated at 'o' and 'x', respectively). In **A**, we simulate an exponentially distributed latent period given by $L = 1$, resulting in a phage infection propagation that outpaces bacteria chemotaxis as the lysis zone keeps expanding outwards (See 2-hour spaced chemotactic front positions overlaid in dark grey over final frame). In **B**, we decouple latent period and resources as $\eta(R) = \frac{\eta}{3}$, where the division by 3 ensures that the maximum lysis rate is the same as in the rest of T7 *vs E. coli* W1485(F8) simulations (see Supplementary Text Section 3). In this case phage infections at the very chemotactic front (overlaid in dark grey every 2 hours), where lysis rate is near its maximum, are transported as in the main figures. On the other hand, phage slowly consume the bacterial lawn behind the transport front, producing a constantly expanding lysis region that differs from the experimentally observed dynamics. All parameters are indicated in Supplementary Text Table S1.

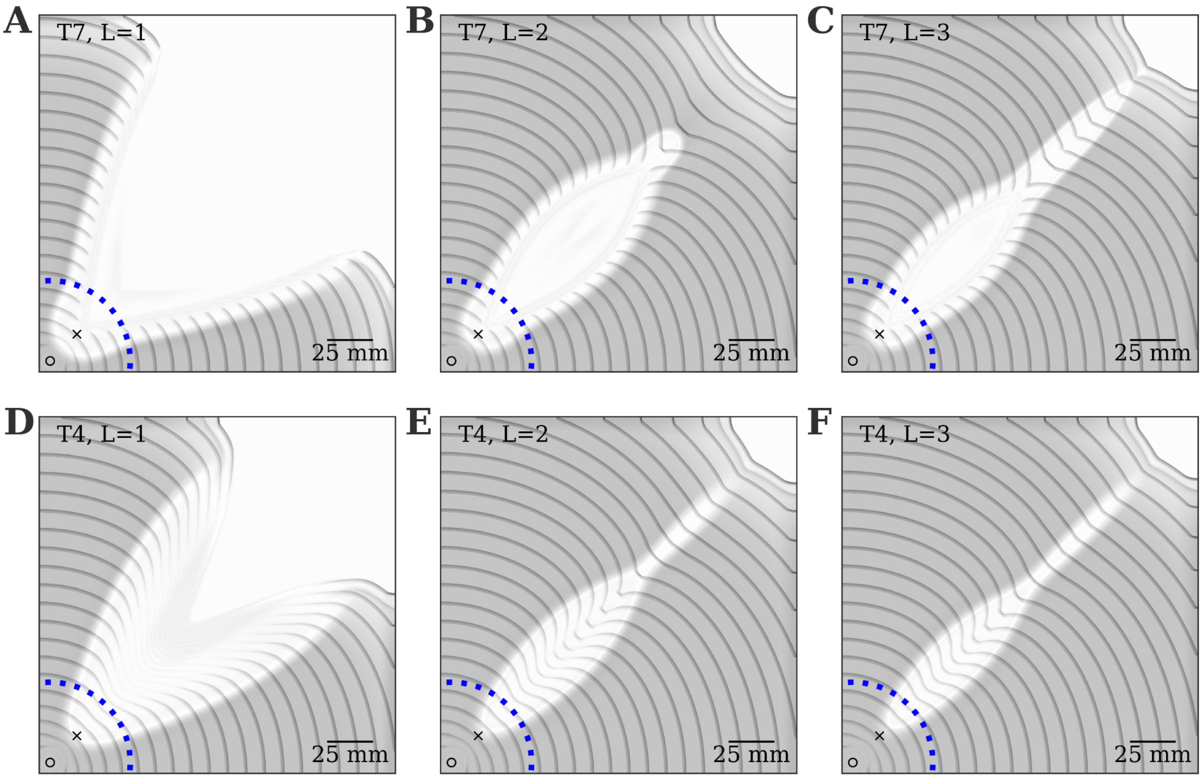


**Supplementary Figure 5.** Simulated dynamics of a PhASER carried by *E. coli* W1485(F8) bacteria (migrating from 'o'), for phages T7 **(A,B,C)** and T4 **(D,E,F)**, inoculated at 'x'. In **A** and **D** we simulate an exponentially distributed latent period (Coefficient of Variation CV=100%) given by $L = 1$, resulting in a phage infection that keeps pace with bacterial chemotaxis and generates tremendous lysis zones that reach the 24cm plate edge without reconverging. We then simulate the model dynamics while increasing $L$ to 2, corresponding to a 70% CV **(B,E)**, and then to $L = 3$, corresponding to a 57% CV **(C,F)**. Final timepoint frames are overlaid with 2 hour-spaced chemotactic front positions (grey curves). Dotted blue lines outline the boundaries of a standard Petri dish with 9 cm diameter. All other parameters are indicated in Supplementary Text Table S1.

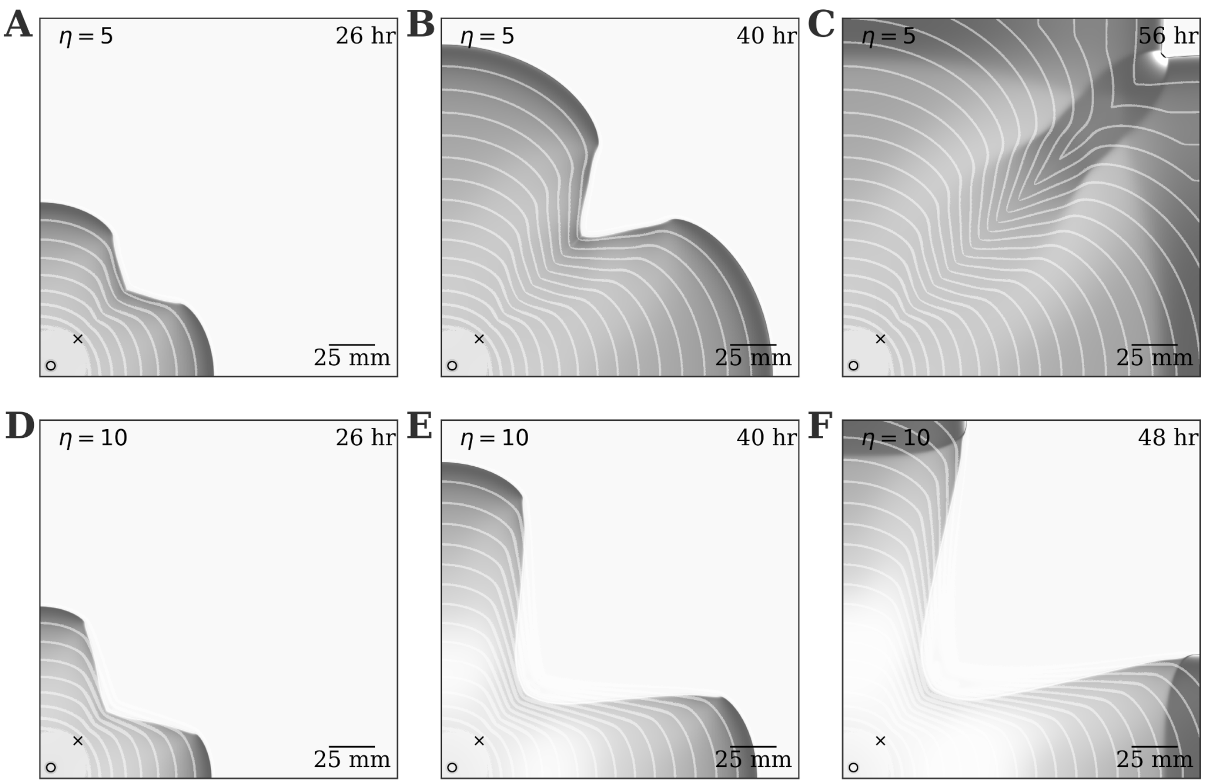


**Supplementary Figure 6.** Simulated dynamics of a T7 PhASER carried by *E. coli* W1485(F8) bacteria that track the main nutrient. $\chi$ is rescaled to $5 \cdot 10^7 \mu m^2$/hrs to keep the range expansion speed comparable with experiments. In **A,B,C** we keep the phage average lysis time fixed to the default T7 $\eta = 5$/hr, showing frames of the same simulation at 26, 40 and 56 hours respectively. In **D,E,F** we increased the lysis time to $\eta = 10$/hr to prevent invasion from the bacteria lawn for example purposes, showing frames of the same simulation at 26, 40 and 48 hours respectively. Final timepoint frames are overlaid with 2 hour-spaced front positions (white curves). All other parameters are indicated in Supplementary Text Table S1, except for $a_-$ and $a_+$, here multiplied by $K/K_2$.

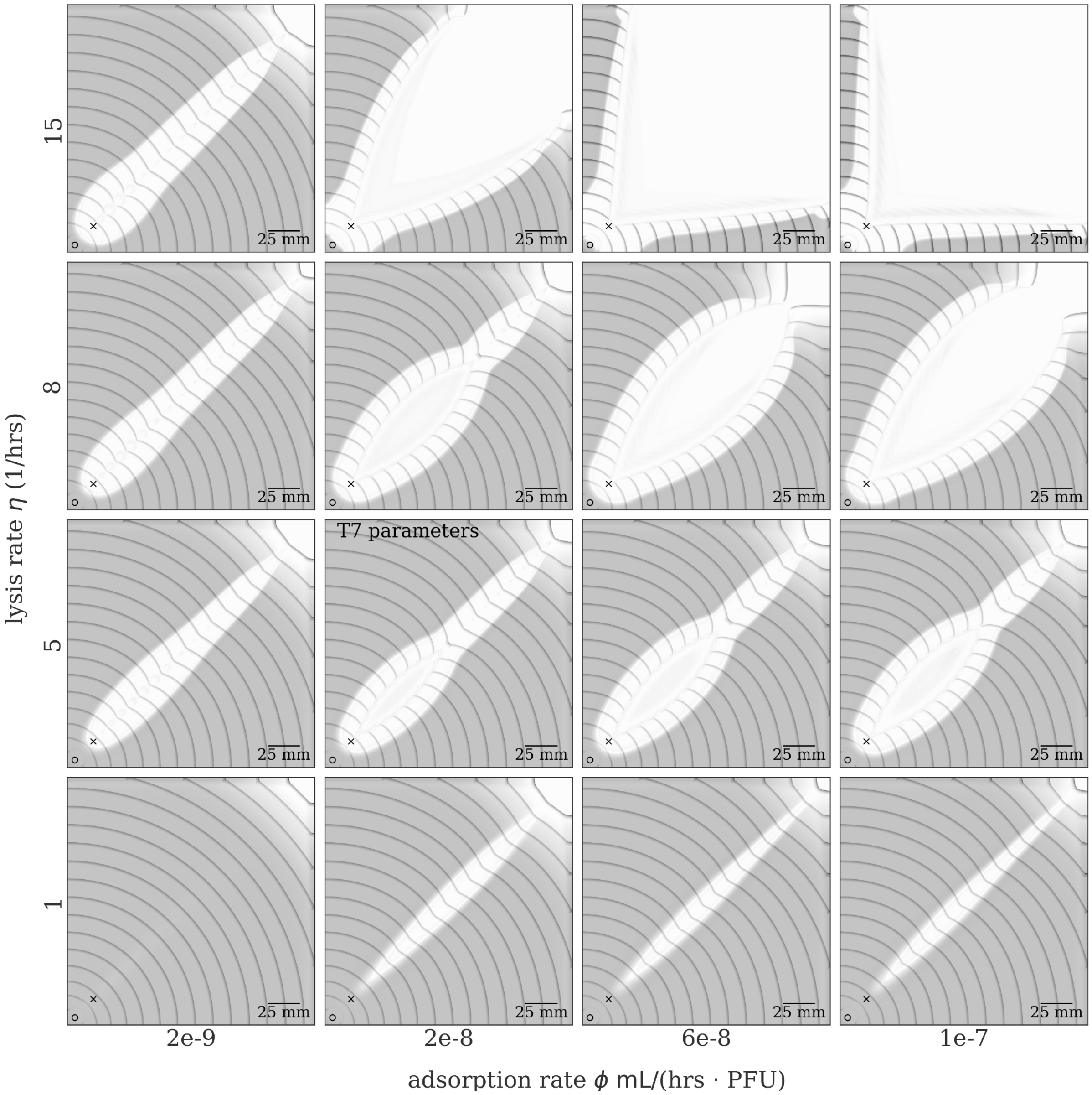


**Supplementary Figure 7.** Simulated dynamics of a PhASER carried by *E. coli* W1485(F9) bacteria when varying the lysis rate $\eta$ between 1 and 15/hr, and the phage adsorption rate $\phi$ between $2 \cdot 10^{-9}$ and $10^{-7}$ mL/(hr PFU). PhASER emergence shows a remarkable robustness to 10-fold variations in viral life traits. For extremely low adsorption rates and lysis rates phage infections are unable to sustain PhASERs. Increasing either adsorption or lysis rates produce stable PhASERs with a wider and wider lysis region, until at very high values the chemotactic fronts at its sides do not reconverge within our experiment's spatial scales. Hosts and phage are inoculated at points 'o' and 'x', respectively. Final timepoint frames are overlaid with 2 hour-spaced front positions (grey curves). All other parameters correspond to those indicated for phage T7 in Supplementary Text Table S1.

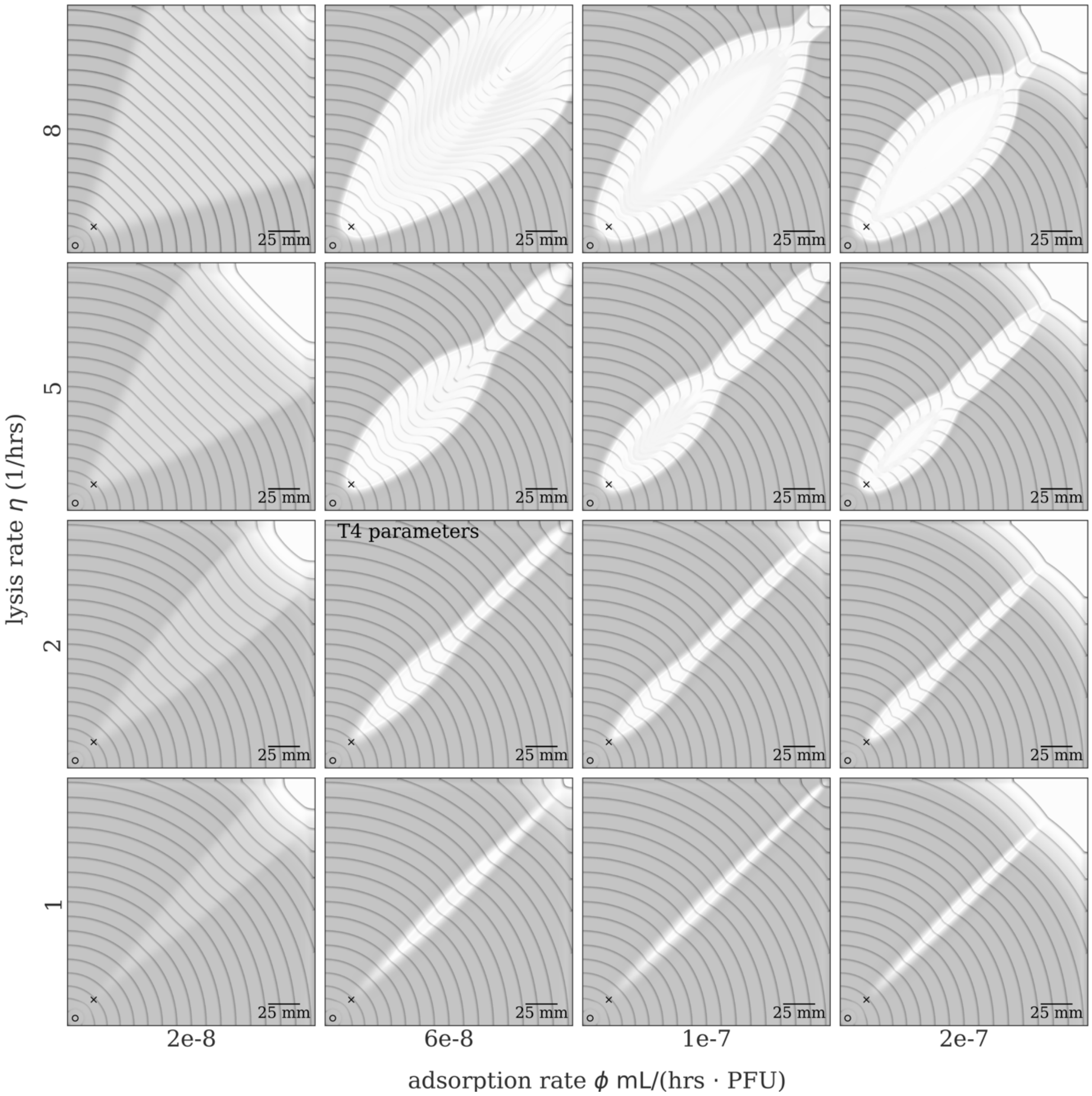


**Supplementary Figure 8.** Simulated dynamics of a PhASER carried by *E. coli* W1485(F9) bacteria when varying the lysis rate $\eta$ between 1 and 8/hr, and the phage adsorption rate $\phi$ between $2 \cdot 10^{-8}$ and $2 \cdot 10^{-7}$ mL/(hr PFU), employing the T4 nonlinear saturation profile $F(P) = \phi \frac{P}{1+P/P_c}$ with $P_c = 10^7$ PFU/mL. PhASER emergence is robust variations in viral life traits spanning 1 order of magnitude. For extremely low adsorption rates phage infections do not drastically reduce bacteria densities at the center of the plate, and the vanishing curvature of the chemotactic front is not pronounced enough to produce the typical PhASER reconvergence within our experimental spatial scales. Increasing either adsorption or lysis rates produces stable narrow PhASERs with residual cells at its center, as observed experimentally for phage T4. Higher adsorption and lysis rates offset the nonlinear phage saturation profile and produce PhASERs that present T7-like features – *i.e.* initial curved fronts divergence and vanishing cell densities inside PhASERs. Hosts and phage are inoculated at points 'o' and 'x', respectively. Final timepoint frames are overlaid with 2 hour-spaced front positions (grey curves). All other parameters correspond to those indicated for phage T4 in Supplementary Text Table S1.

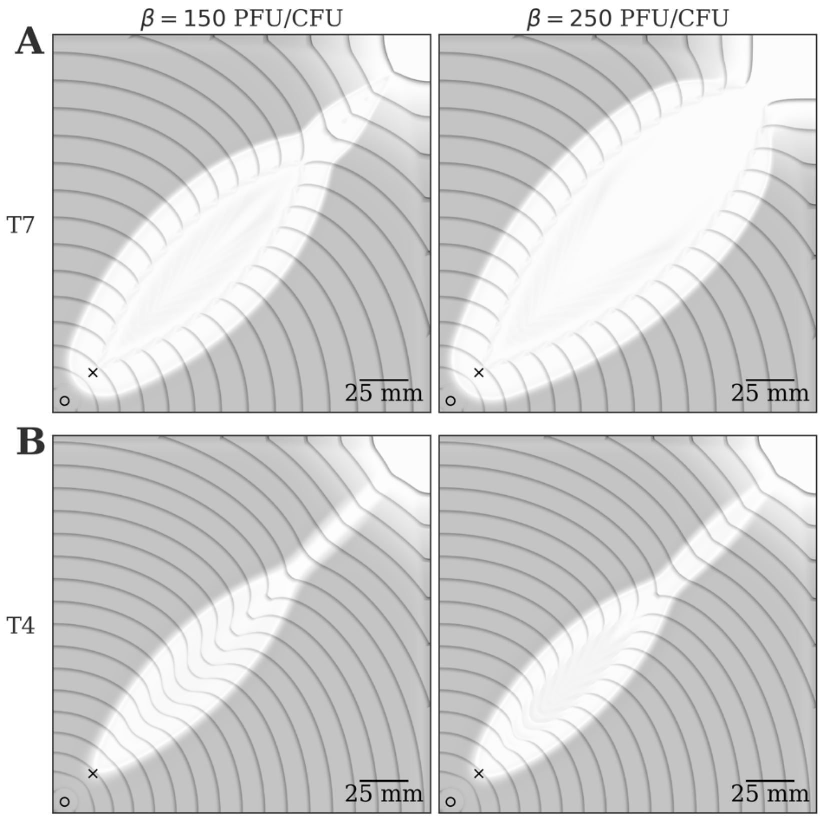


**Supplementary Figure 9.** Simulated dynamics of a PhASER carried by *E. coli* W1485(F9) bacteria when increasing the burst size $\beta$ to 150 and 250 PFU/CFU, for phage T7 **(A)**, and T4 **(B)**. With $\beta =$ 150 PFU/CFU we recover qualitatively similar PhASER dynamics as in Figure 2 in the main text. A higher $\beta =$ 250 PFU/CFU yields T7 lysis regions that do not reconverge within our experiment's spatial scales, while T4 dynamics start presenting quantitative PhASER features typically associated with phage T7 – *i.e.* initial curved fronts divergence and vanishing cell densities inside PhASERs. Hosts and phage are inoculated at points 'o' and 'x', respectively. Final timepoint frames are overlaid with 2 hour-spaced front positions (grey curves). All other phage-dependent parameters correspond to those indicated in Supplementary Text Table S1.

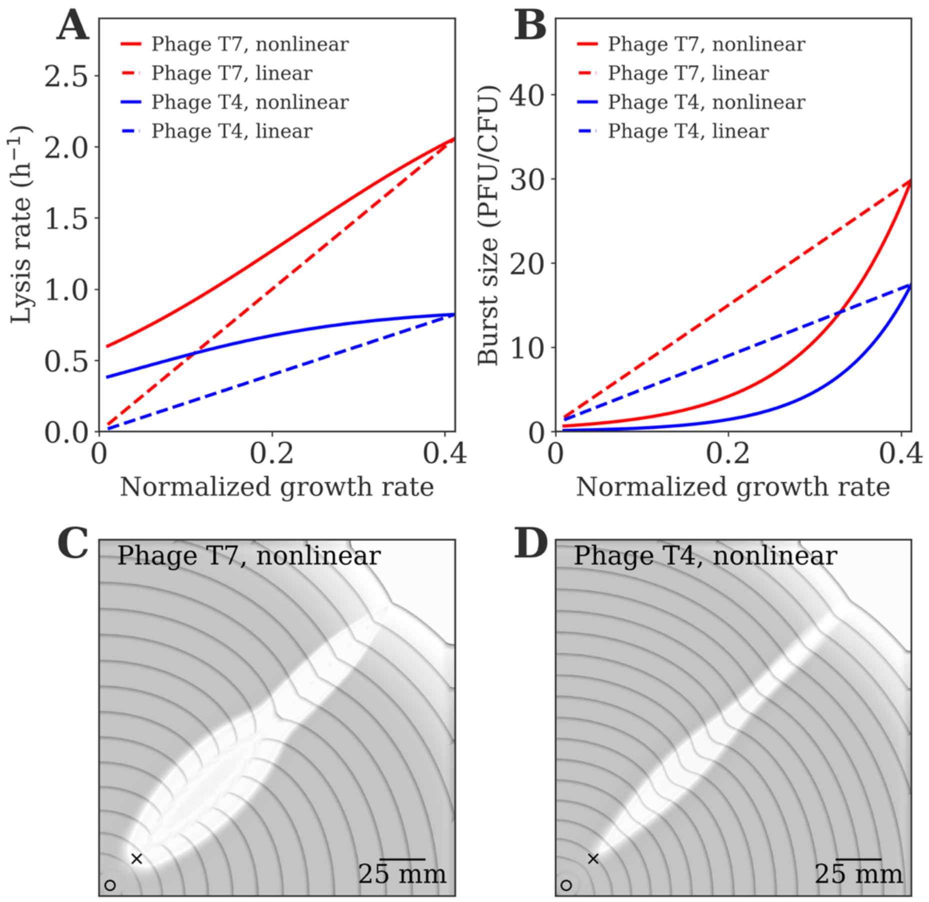


**Supplementary Figure 10.** Simulated dynamics of a PhASER model with nonlinear dependence of phage lysis rate and burst size with bacteria growth rate. Panels **A,B** compare the linear functions employed in the main text (dashed lines) for phage T7 (red) and T4 (blue), with the nonlinear lysis rate and burst size functions specified in equations S1 and S2 in the Supplementary text (full lines). These functions were adapted from empirically-informed phage traits dependence on hosts growth rate[47], details in the Supplementary text. The x-axis spans the normalized bacteria growth rate range realized in the simulations based on the resources dynamics parameters – *i.e.* $\left[0, \frac{R_0}{R_0+K}\right]$, see Supplementary Text. Panels **C,D** show spatial simulations for phage T7 and T4 respectively with nonlinear phage traits, resulting in the formation of stable PhASERs. All other parameters are kept the same as in Figure 2 of the main text, as indicated in Table S1 for *E. coli* strain W1485(F9).

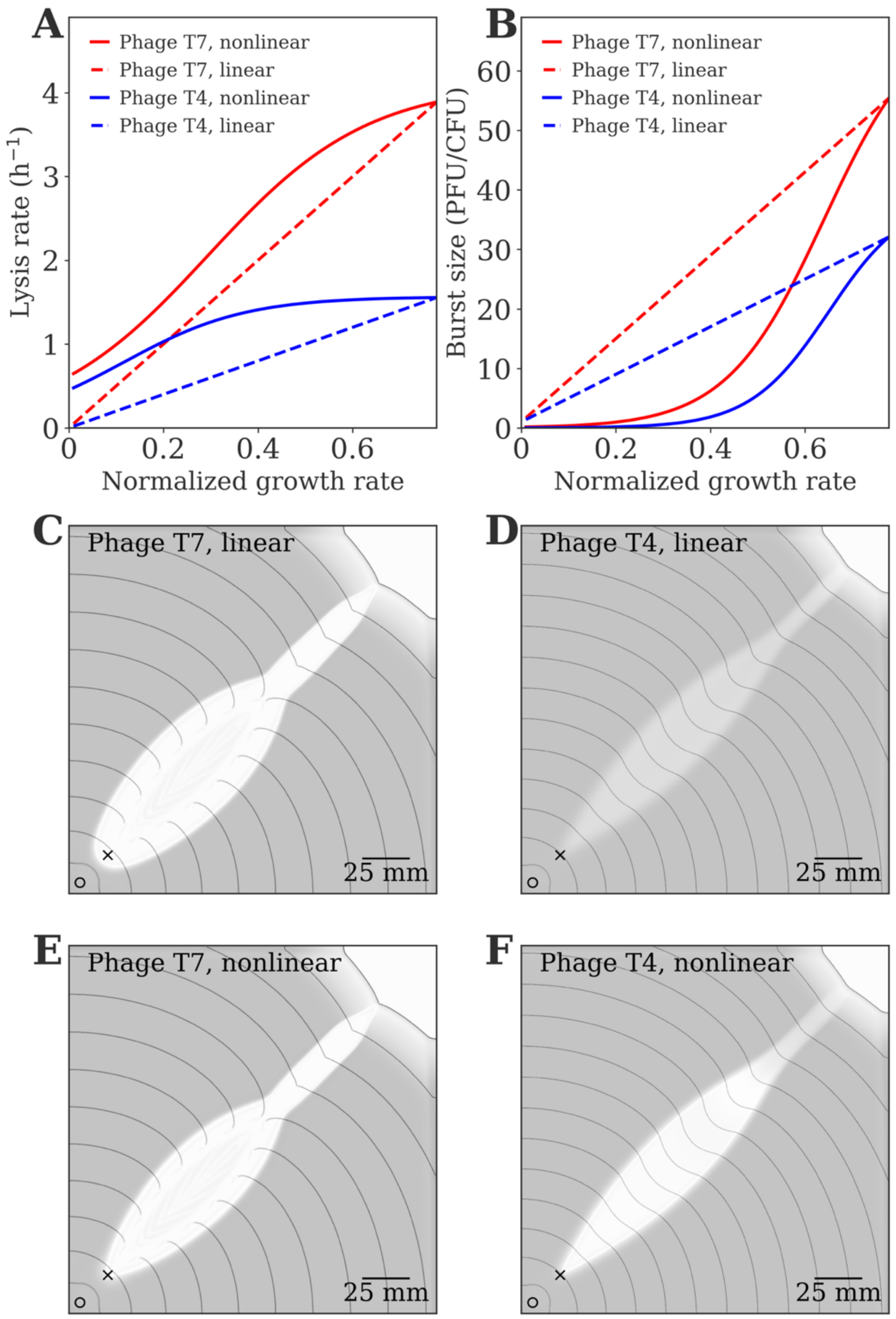


**Supplementary Figure 11.** Simulated dynamics of a PhASER model with a linear versus nonlinear dependence of phage lysis rate and burst size with bacteria growth rate, upon modifying the Monod constant parameter $K$ from $5 \cdot 10^9$ to $10^9$ CFU/mL – indicated in units of host cell concentrations as explained in the Supplementary text. Panels **A,B** compare the linear lysis rate and burst size functions (dashed lines) for phage T7 (red) and T4 (blue), with the nonlinear ones specified in equations S1 and S2 in the Supplementary text (full lines). The x-axis spans the normalized bacteria growth rate range realized in the simulations based on the resources dynamics parameters – *i.e.* $\left[0, \frac{R_0}{R_0+K}\right]$, which is almost doubled compared to the baseline simulations, see Supplementary Text. Panels **C,D** show spatial simulations for phage T7 and T4 respectively with linear phage traits, while panels **E,F** show the model outcomes with the nonlinear functions plotted in **A,B**. Despite quantitative differences, all of these simulations resulted in the formation of stable PhASERs. All other parameters are kept the same as in Figure 2 of the main text, as indicated in Table S1 for *E. coli* strain W1485(F9).

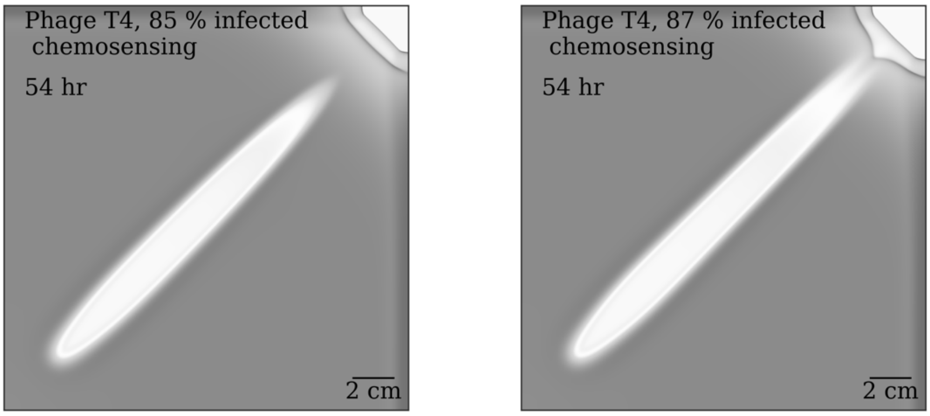


**Supplementary Figure 12.** Simulated dynamics of a T4 *vs E. coli* W1485(F8) PhASER model where the parameter governing infected cells' responsiveness to chemoattractant gradients is modulated as $\tilde{\chi} = \alpha\chi$ with $\alpha$ between 0 and 1. The stable PhASER propagation towards the top-right corner is lost when $\alpha$ is decreased from 0.87 to 0.85. All parameters are indicated in Supplementary Text Table S1.

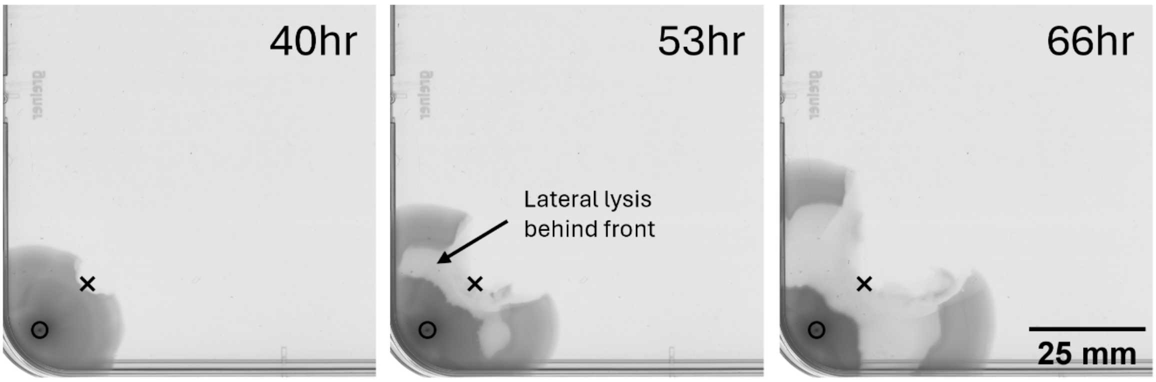


**Supplementary Figure 13.** Deviation from focused infection transport at low migration speeds in higher concentration swim agar. A colony (dark grey) of *E. coli* W1485(F8) grows and migrates at ~ 0.7mm/hr through an 0.4% agar LB swim assay from location ∘, encounters T7 phage at location ×, and displays a rapid lateral spread of lysis behind the migrating front. The timeseries is labelled with hours post inoculation.

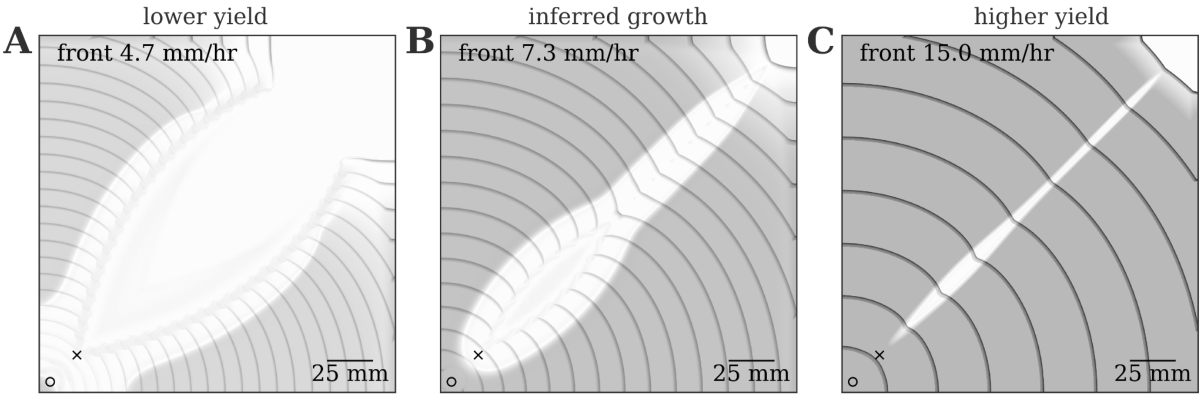


**Supplementary Figure 14.** Simulated dynamics of a PhASER carried by *E. coli* W1485(F9) bacteria (migrating from 'o'), for phage T7, inoculated at 'x'. In these simulations we modulated yield, *i.e.* the biomass conversion factor between nutrients and cells (as detailed in Supplementary Text), producing a growth rate either half **(A)** or four times **(C)** that originally inferred for this bacterial strain **(B)**. Note that, as expected, the population-mediated range expansion speed depends on the growth rate. Time frames are overlaid with 2 hour-spaced front positions (grey curves). All other parameters are indicated in Supplementary Text Table S1.

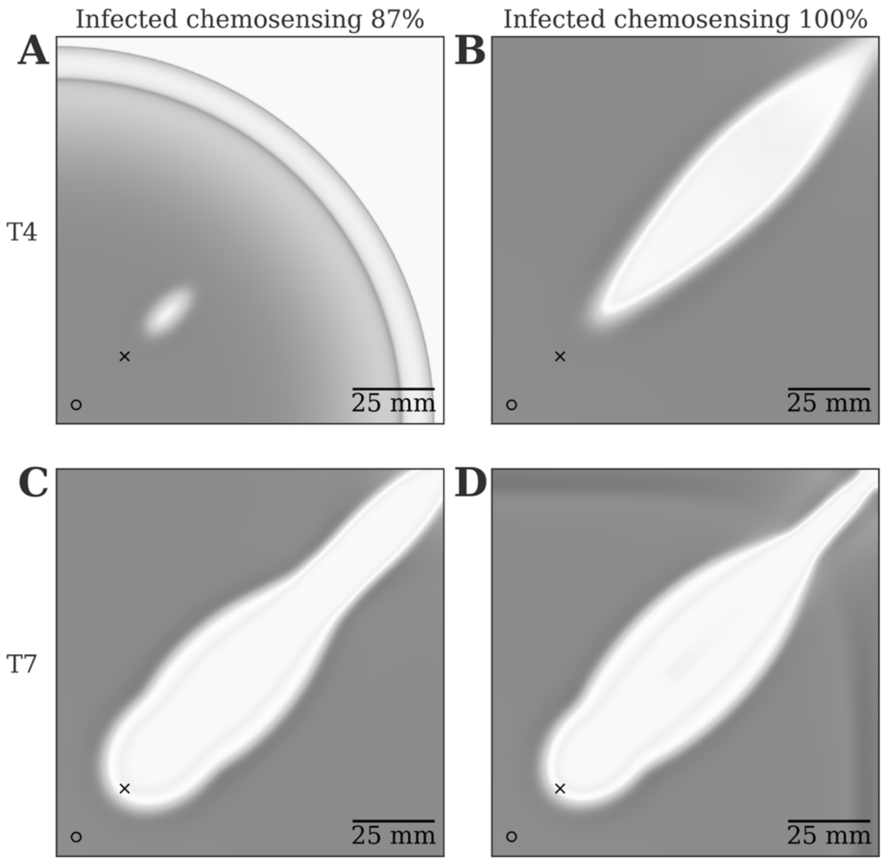


**Supplementary Figure 15.** Simulated dynamics of a T4 **(A,B)** and T7 **(C,D)** *vs E. coli* W1485(F8) PhASER model, where the parameter governing infected cells' responsiveness to chemoattractant gradients is modulated as $\tilde{\chi} = \alpha\chi$ with $\alpha = 0.87$ **(A,C)** and $\alpha = 1$ **(B,D)**. The simulations are initialized with a very low phage inoculum of $5 \cdot 10^5$ PFU/mL, leading to an estimated phage density of $4.7\ \Theta$ just before phage and bacteria come in contact. This choice corresponds to the phage inoculum in the second column of Figure 4E, with those two plots replotted here as panels **A** and **D** for comparison. For phage T4, reducing $\alpha$ leads to the dependence of stable PhASER propagation on phage inoculum density (**A** and Figure 4E), which is not realized with $\alpha = 1$ **(B)**. For phage T7, an equivalent reduction in $\alpha$ leads to a stable PhASER even at such low inoculum **(C)**, and therefore phage inoculum density does not affect PhASER formation like for $\alpha = 1$ (**D** and Figure 4E). All other phage-dependent parameters are indicated in Supplementary Text Table S1.

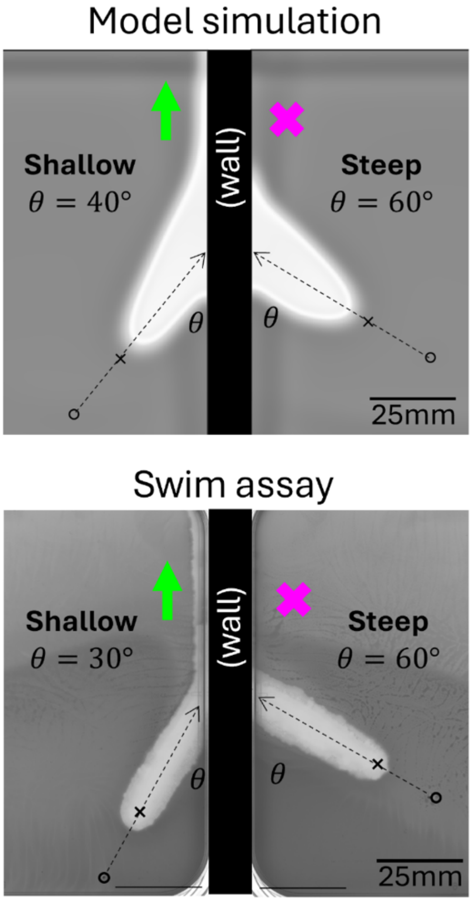


**Supplementary Figure 16.** PhASERs intersecting solid walls terminate or turn and propagate stably along them, depending on angle. Model simulations (top) and experimental swim assays (bottom) in which a bacterial host (*E. coli* W1485(F8) or parameters fit to same) grows and chemotaxes from an inoculation point ∘, to eventually cover a swim assay plate (dark grey growth). On encountering a spot of lytic phage (T4 or parameters fit to same) inoculated 20mm away at ×, a PhASER (cleared zone) forms in the migrating bacterial front and propagates straight (dashed arrow) until intersecting a solid wall. On the left, PhASERs intersecting the wall at shallow angles of roughly half that of the merging PhASERs in Figure 5A, in the simulation ($40°$) or swim assay ($30°$) turn and continue to propagate with the migrating host front along the wall (green arrows). PhASERs intersecting the wall at a steep angle (60°, half of merger angles in Figure 5B) lose the advancing front on collision and terminate (magenta crosses).

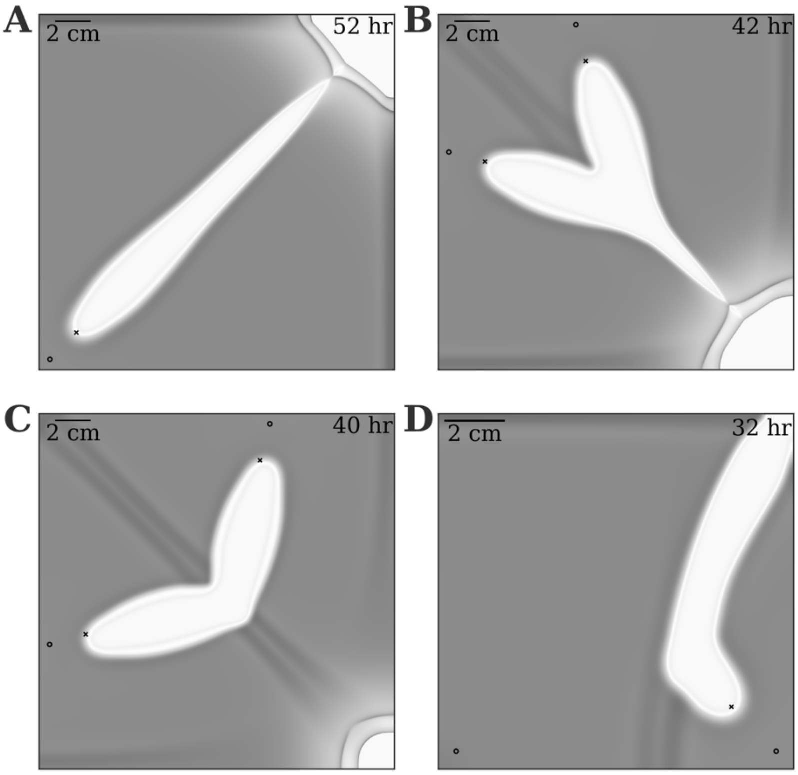


**Supplementary Figure 17.** Simulated dynamics of a T4 PhASER model with a linear adsorption profile $F(P) = \phi P$, transported by *E. coli* W1485(F8). The simpler adsorption profile maintains: **(A)** PhASER formation and stability, **(B)** robustness to collisions with other communities at 60 degrees , **(C)** PhASER termination upon collision at 120 degrees , and **(D)** stability to ecological perturbations challenging PhASERs with an *E. coli* W1485(F10) colony (inoculated at bottom-left) expanding at a much higher speed than W1485(F8) at bottom-right . Host and phage are inoculated at locations marked 'o' and 'x', respectively. All parameters are indicated in Supplementary Text Table S1, except for $\phi = 2 \cdot 10^{-9}$ mL/(hrs PFU), $\eta = 5$ hrs$^{-1}$ and $\beta = 50$.

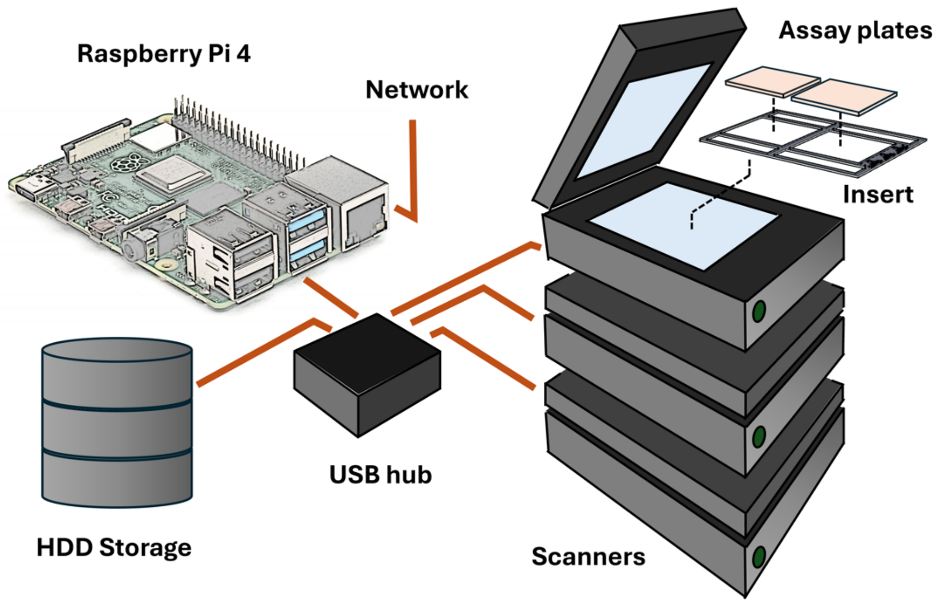


**Supplementary Figure 18.** Raspberry Pi-connected scanner cluster layout. Independent Raspberry Pi-controlled scanner timelapse modules are used to allow flexibility in running parallel experiments. Modules are composed of a Raspberry Pi 4 computer connected via USB hub to groups of (here, three Epson V800/850) scanners, and a local storage drive. The Raspberry Pi computers are run headless over the local network using VNC. A custom python script (Supplementary Method 1) is used to specify timelapse and image parameters and schedule scans and uses the SANE (http://www.sane-project.org/) interface to the scanners. Scanners are placed within temperature-controlled chambers. Custom 3D-printed inserts (Supplementary Method 2) are used to repeatably locate assay plates in the transmitted light imaging area of scanner platens.

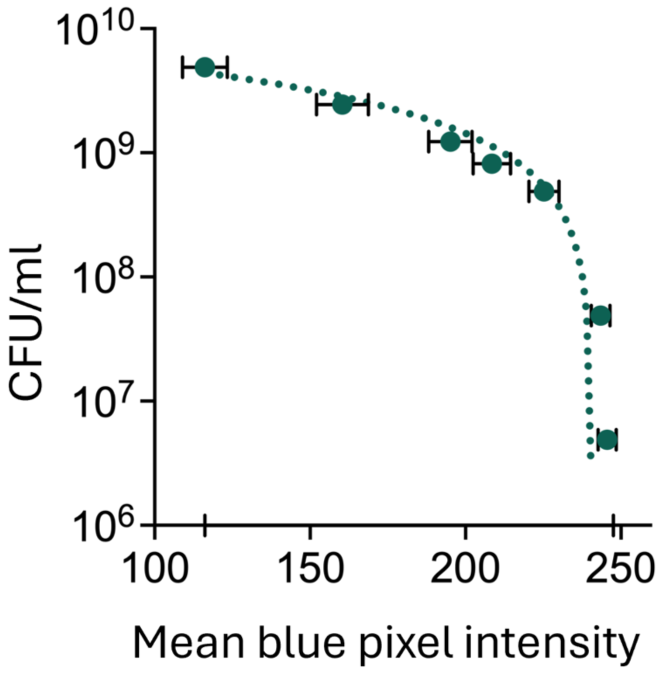


**Supplementary Figure 19.** Calibration curve (dashed, linear regression) used to map blue channel pixel intensity values to bacterial concentration (colony forming units (CFU) per ml) in scanned swim plate images (see Methods). 8-bit pixel intensities range from 0 to 255. Error bars represent standard deviation in pixel value across the plate area.

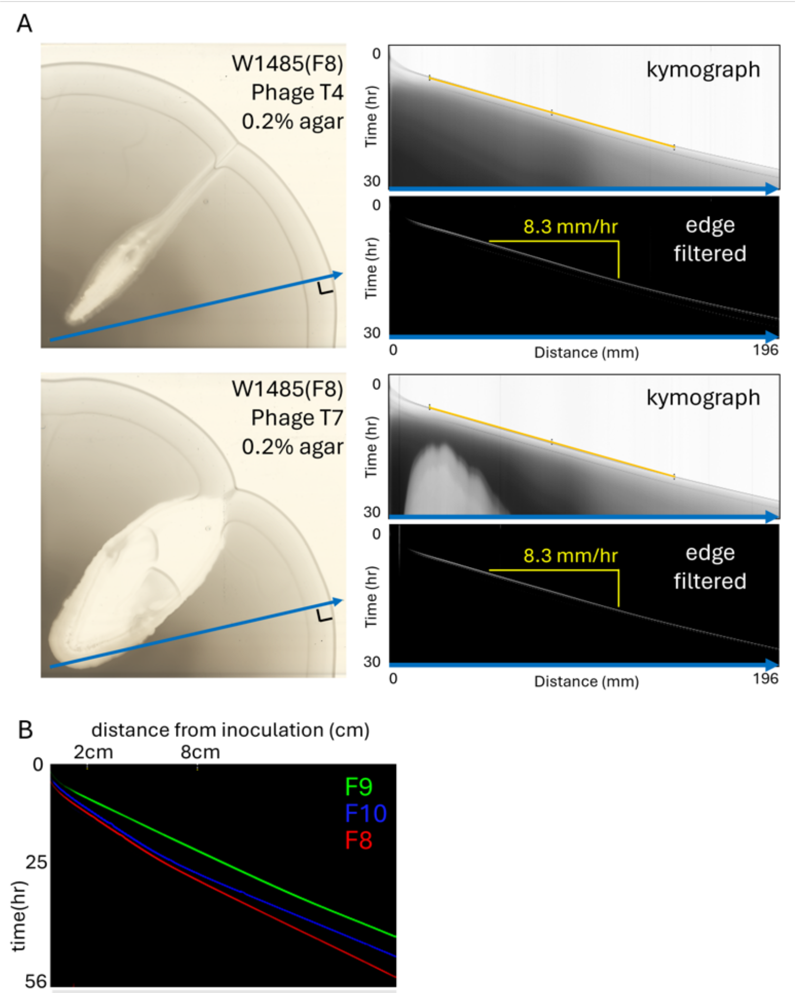


| | 2cm + 10hr | 8cm + 10hr |
|---|---|---|
| W1485(F8) | 5.5 mm/hr | 6.9 mm/hr |
| W1485(F9) | 7.1 mm/hr | 7.4 mm/hr |
| W1485(F10) | 5.8mm/hr | 8.7 mm/hr |

**Supplementary Figure 20.** Determining chemotactic range expansion rate using kymographs. **A.** Linear regions of interest (ROIs, blue arrows on scanned images of assay plates from figure 3A) that are orthogonal to the migrating fronts throughout the timelapse sequence are specified, often extending from bacterial inoculation points. Kymographs, pixel intensity maps along the linear ROIs through time, are extracted from the full timelapse sequence (FIJI Multi Kymograph plugin) and displayed for the phage T4 (top) and T7 (bottom) assays. The orientation of the line profile is indicated by the blue arrow at the base (final timepoint) of the kymographs. Straight diagonal edges in these kymographs (overlaid by the orange line and highlighted using a sobel edge filter) indicate constant speed of advancement of the migrating front along the blue line. The reciprocal slope of this edge is the speed of the migrating front along the linear ROI. Speed during PhASER initiation is estimated by fitting front migration for 20 hours from phage inoculum contact. **B.** We observe that our *E. coli* assay strains with two chemotactic fronts (W1485(F8), W1485(F10)) tend to uniformly adjust to quicker migration speeds after ~20 hrs growth. Here, for migration in LB swim medium with 0.25% agar, the speed change is evident in the bent red and blue edge-filtered first migration fronts in the kymograph, and differences in each strain's measured front speed over the 10 hours after reaching 2cm and 8cm from inoculation. The speed adjustment is weaker in the W1485(F9) mutant strain which has a single dominant migration front (see straighter green front trajectory and more slight speed change).

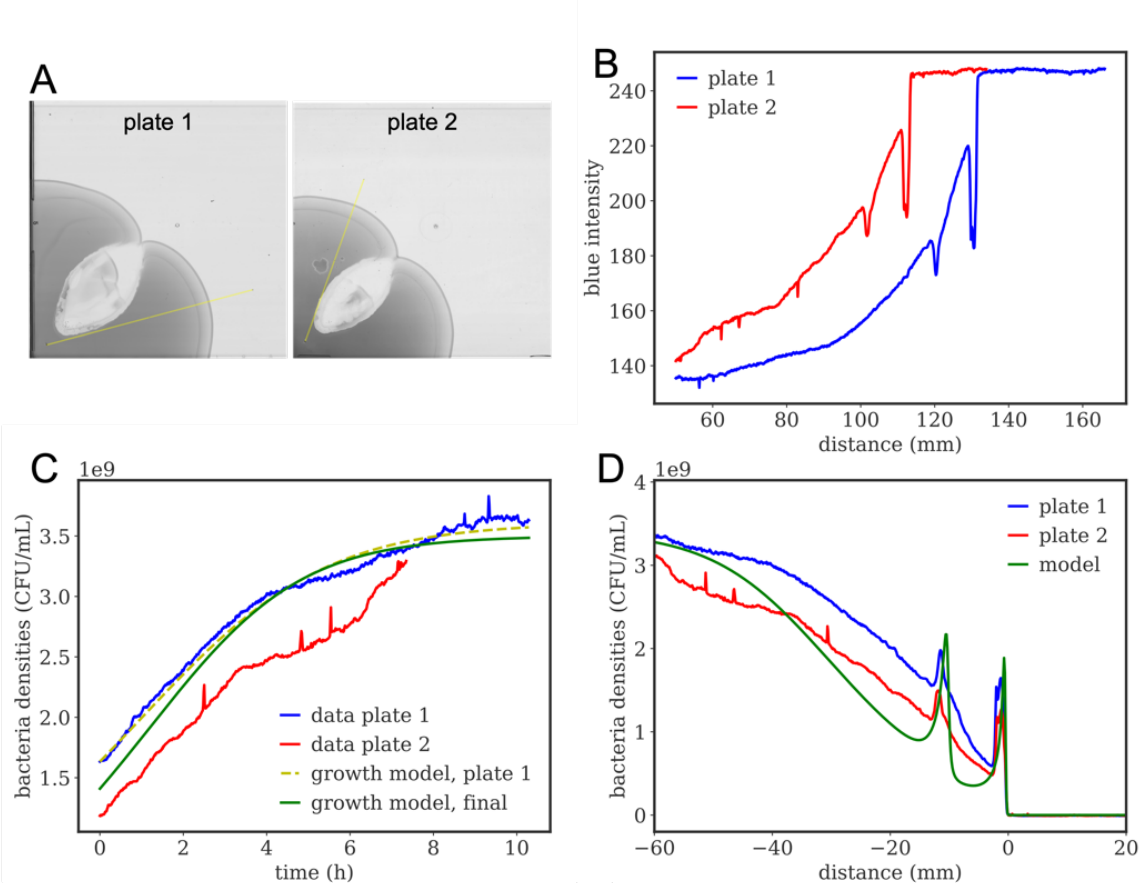


**Supplementary Figure 21.** Fitting pipeline to fix *E. coli* W1485(F8) bacteria life trait parameters. First, we select two plates with an expanding W1485(F8) colony, slicing the bacteria optical density along a radius parallel to the range expansion direction, in a region unaffected by phage lysis **(A)**. Then we record the blue channel intensity along the selected lines **(B)**, which is transformed to bacteria densities (CFU/mL) according to the calibration data in Supplementary Figure 9. Using the wave properties of the range expansion progressing at speed $v$, we transform the radial density profile measured a distance $\Delta$ behind the front – discarding the profile up to the last front if more than one is present – to growth that has been taking place for a time $t = \frac{\Delta}{v}$ since the front was at that location seeding cells behind it **(C)**. We use this indirectly derived growth curve to infer the parameters of a simple Monod growth model (see Supplementary Text Section 2 for details), where the yellow dashed line shows the growth model outcome with the parameters inferred on the plate 1 profile, while the green line shows the growth model dynamics with common parameters reproducing features from both plates (in this example, the fact that plate 2 initially has a higher growth rate, see Supplementary Text Section 2). Finally, we use these growth parameters in the full PDE model introduced in Box 1 to fit two parameters controlling chemical sensing and aspartate consumption rate matching the experimental radial density profile **(D)**. Bacteria responsiveness to sensed gradients is fixed to match $v$. Other W1485(F8) trait values are fixed from literature (see Supplementary Text Section 2).

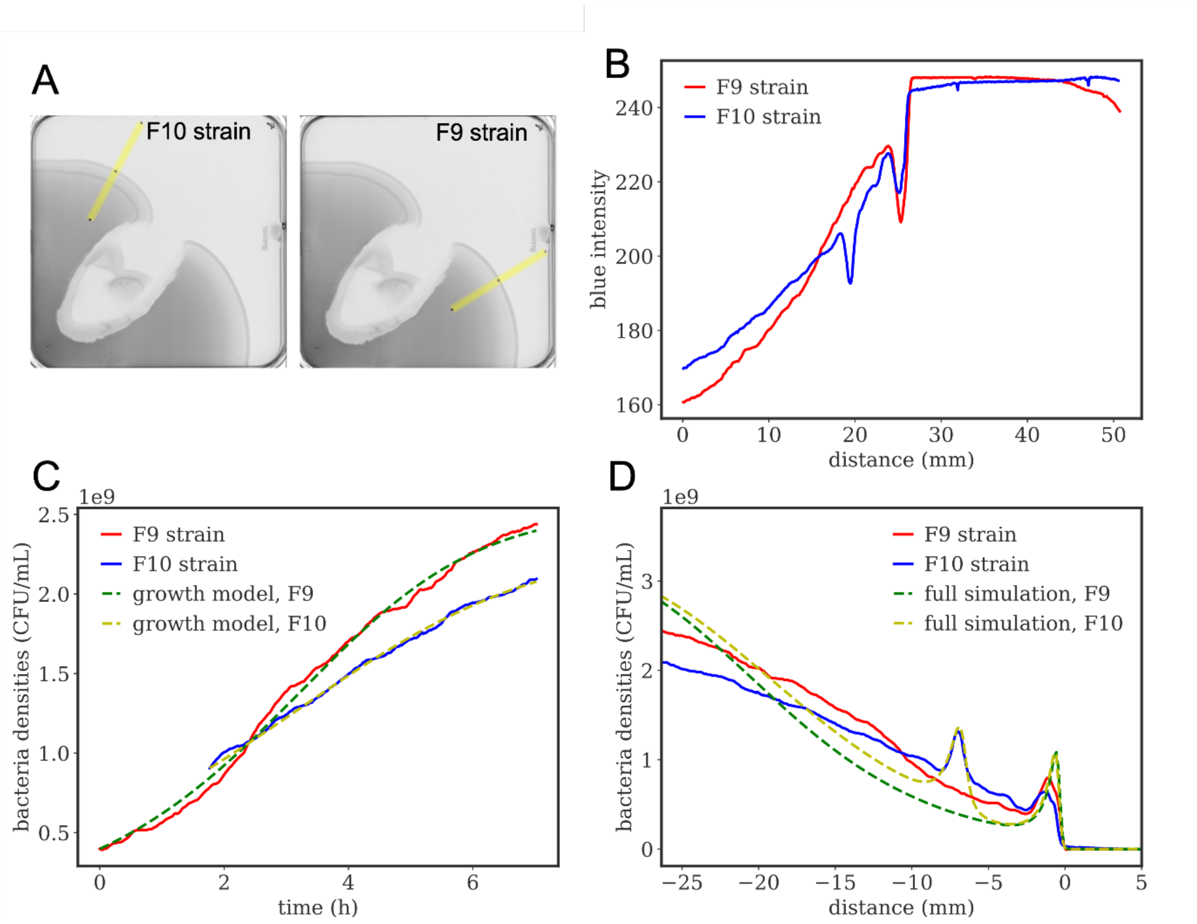


**Supplementary Figure 22.** Fitting pipeline to fix *E. coli* W1485(F9) and W1485(F10) strain life trait parameters. First, we select a plate with an expanding colony featuring *E. coli* W1485(F9) on the right of the plate and *E.coli* W1485(F10) on the left, slicing the bacteria optical density along a radius parallel to the range expansion direction, in a region unaffected by phage lysis **(A)**. For each strain we record the blue channel intensity along the selected lines **(B)**, which is transformed to bacteria densities (CFU/mL) according to the calibration data in Supplementary Figure 9. Then, using the wave properties of the range expansion progressing at speed $v$, we transform the radial density profile measured a distance $\Delta$ behind the front – discarding the profile up to the last front if more than one is present – to growth that has been taking place for a time $t = \frac{\Delta}{v}$ since the front was at that location seeding cells behind it **(C)**. We use this indirectly derived growth curve to infer the parameters of a simple Monod growth model (see Supplementary Text Section 2 for details), where the yellow dashed line shows the growth model outcome with the parameters inferred for strain W1485(F10) (blue), while the green dashed line shows the growth model dynamics with parameters inferred for strain W1485(F9). To avoid overfitting we choose common parameters for these two strains, compatible with experimental variation, to simulate the full PDE model introduced in Box 1 (details in Supplementary Text Section 2). Assuming the only difference between the two strains is W1485(F9) insensitivity to one of the two attractants, we fit a few sensing and uptake parameters (see Supplementary Text Section 2) to match the radial density profiles measured experimentally for each strain **(D)**.
