## Supplementary Text for "Stable coexistence and transport of lytic phage infections with migrating bacterial hosts"

### 1. NUMERICAL IMPLEMENTATION

We simulate the Partial Differential Equation model introduced in Box 1 with a finite differences scheme. We introduce a 2D grid with 100 $\mu$m spacing, where population concentrations are tracked in time, mimicking the experimentally measured concentrations averaged in the vertical direction. We assign a virtual (*i.e.*, not explicitly simulated) height of 0.5cm to the grid, compatible with the soft agar layer in our experimental setup. The volume of each lattice site corresponds to $0.5 \cdot 10^{-4}$ mL. The size of the grid is either set to 12 or 20 cm, to match the corresponding agar plate format used in the experiments.

The spatial derivatives $\frac{\partial}{\partial x}, \frac{\partial}{\partial y}$ present in the diffusion and advection terms are discretized on the square lattice. We use the following stencil kernel to discretize the Laplacian operator ($\nabla^2 = \frac{\partial}{\partial x}^2 + \frac{\partial}{\partial y}^2$) in the diffusion terms:

$$\text{Kernel} = \frac{1}{\Delta x^2}\begin{pmatrix} 0.25 & 0.5 & 0.25 \\ 0.5 & -3 & 0.5 \\ 0.25 & 0.5 & 0.25 \end{pmatrix}$$

We convolve this kernel with the state fields. For instance, at time $t$, the diffusion term of resources $D_R \nabla^2 R$ at a site (x,y) can be obtained by discretizing spatial derivatives to first order with respect to first- and second-neighbor sites:

$$\begin{aligned}
D_R \nabla^2 R(t,\mathbf{x})|_{(x,y)} &= \frac{D_R}{\Delta x^2}\begin{pmatrix} 0.25 & 0.5 & 0.25 \\ 0.5 & -3 & 0.5 \\ 0.25 & 0.5 & 0.25 \end{pmatrix} * R(t,\mathbf{x})|_{(x,y)} \\
&= \frac{D_R}{\Delta x^2}\begin{pmatrix} 0.25 & 0.5 & 0.25 \\ 0.5 & -3 & 0.5 \\ 0.25 & 0.5 & 0.25 \end{pmatrix} \cdot \begin{pmatrix} R(t,x-\Delta x,y+\Delta y) & R(t,x,y+\Delta y) & R(t,x+\Delta x,y+\Delta y)) \\ R(t,x-\Delta x,y) & R(t,x,y) & R(t,x+\Delta x,y) \\ R(t,x-\Delta x,y-\Delta y) & R(t,x,y-\Delta y) & R(t,x+\Delta x,y-\Delta y) \end{pmatrix} \\
&= \frac{D_R}{\Delta x^2}\Big( -3R(t,x,y) + 0.5\Big(R(t,x+\Delta x,y) + R(t,x-\Delta x,y) + R(t,x,y+\Delta y) + R(t,x,y-\Delta y)\Big) + \\
&+ 0.25\Big(R(t,x+\Delta x,y+\Delta y) + R(t,x-\Delta x,y+\Delta y) + R(t,x+\Delta x,y-\Delta y + R(t,x-\Delta x,y-\Delta y)\Big)\Big),
\end{aligned}$$

where $*$ denotes a convolution operation while $\cdot$ denotes the inner product. Diffusion of viruses, chemoattractants and bacteria is given by the same equation accounting for their specific diffusion constant, noting that the bacteria diffusion term represents the random drift component of an active run-and tumble motion rather than passive Brownian motion.

The divergence of the velocity field determining chemotaxis takes the general form $\nabla \cdot f(\mathbf{x})\nabla g(\mathbf{x}) = \frac{\partial}{\partial x}\left[f(\mathbf{x})\frac{\partial}{\partial x}g(\mathbf{x})\right] + \frac{\partial}{\partial y}\left[f(\mathbf{x})\frac{\partial}{\partial y}g(\mathbf{x})\right]$ where the function $f$ stands for the corresponding bacteria class density, while the function $g$ stands for the chemosensing response $\chi \ln\left(\frac{1+a_n/a_-}{1+a_n/a_+}\right)$. We evaluate this differential operator on the discrete grid through a central differencing scheme. For example to evaluate $\frac{\partial}{\partial x}\left[f(\mathbf{x})\frac{\partial}{\partial x}g(\mathbf{x})\right]$ at a site (x,y), we interpolate $f$ on the faces of the lattice site as $f(x-\frac{\Delta x}{2},y) = \frac{f(x,y)+f(x-\Delta x,y)}{2}$ and $f(x+\frac{\Delta x}{2},y) = \frac{f(x,y)+f(x+\Delta x,y)}{2}$. This allows us to evaluate the flux $f(\mathbf{x})\frac{\partial}{\partial x}g(\mathbf{x})$ at the site face $(x+\frac{\Delta x}{2},y)$ as $f(x+\frac{\Delta x}{2},y)\frac{g(x+\Delta x,y)-g(x,y)}{\Delta x}$. Finally, differentiating the fluxes on the site faces, we have:

$$\begin{aligned}
\frac{\partial}{\partial x}\left[f(\mathbf{x})\frac{\partial}{\partial x}g(\mathbf{x})\right]_{(x,y)} =& f(x+\frac{\Delta x}{2},y)\frac{g(x+\Delta x,y)-g(x,y)}{\Delta x^2} - f(x-\frac{\Delta x}{2},y)\frac{g(x,y)-g(x-\Delta x,y)}{\Delta x^2} \\
=& \frac{1}{2\Delta x^2}[f(x+\Delta x,y)g(x+\Delta x,y) + f(x-\Delta x,y)g(x-\Delta x,y) - 2f(x,y)g(x,y) \\
&+ f(x,y)g(x+\Delta x,y) + f(x,y)g(x-\Delta x,y) - f(x+\Delta x,y)g(x,y) - f(x-\Delta x,y)g(x,y)].
\end{aligned}$$

Generalizing to both dimensions through the whole grid we have:

$$\nabla \cdot f(\mathbf{x})\nabla g(\mathbf{x}) = \frac{1}{2\Delta x^2}\left[\begin{pmatrix} 0 & 1 & 0 \\ 1 & -4 & 1 \\ 0 & 1 & 0 \end{pmatrix} * f(\mathbf{x})g(\mathbf{x}) + f(\mathbf{x})\begin{pmatrix} 0 & 1 & 0 \\ 1 & 0 & 1 \\ 0 & 1 & 0 \end{pmatrix} * g(\mathbf{x}) - g(\mathbf{x})\begin{pmatrix} 0 & 1 & 0 \\ 1 & 0 & 1 \\ 0 & 1 & 0 \end{pmatrix} * f(\mathbf{x})\right].$$

When calculating spatial derivatives at the boundaries of the grid, we implement reflecting boundary conditions by adding a virtual site beyond the grid mirroring all population densities in the closest site of the real lattice, ensuring that all fluxes across the boundaries are 0. For example, at the right boundary $x = \Omega$, when calculating the convolutions with the diffusion and advection kernels for resources $R$, we set $R(t, \Omega + \Delta x, y) = R(t, \Omega, y)$. Together with the site-specific reaction terms, the spatial terms complete the computation of time derivatives given the state variables at a given time.

As the time derivatives are estimated, the system state is updated to the following time step through an explicit Euler scheme. The time step is changed adaptively throughout the simulation to increase the precision of the numerical integration. At each time iteration we check whether densities are updated to unfeasible negative values by fast reactions. In practice, we flag resource densities below $-K/1000$, attractant densities below $-3a_-$ and phage densities below $-\Theta/10$. If any of these conditions is met, the last iteration is disregarded and the time step is divided by 1.5 before repeating. Otherwise, negative concentrations are set to 0 and the system is updated to the next step, while the time step is multiplied by 1.3 up to a maximum of $5 \cdot 10^{-4}$ hr. The population densities are saved every 2 hours. When plotting the intensities profiles we convert CFU/mL to pixel intensities according to the calibration data in Supplementary Fig. 19.

Simulations are initialized with a constant level of resources and attractants, $R_0$ and $a_0$ respectively (specified and motivated below). Bacteria are initialized as a 2D Gaussian profile centered around the initial position $(x, y) = (6, 6)$mm, where the maximum density is set to $8 \cdot 10^8$ CFU/mL. The standard deviation of the initial profile is set to 2 mm. Similarly, phage are initialized as a Gaussian profile centered around $(21, 21)$mm with standard deviation 0.2mm and maximum value $10^9$ PFU/mL. These initial conditions ensure that the peaks of bacteria and viruses distributions are at $1.5\sqrt{2} \sim 2$ cm similar to the inoculi distance in the experiments. At the beginning of the simulation there are no infected cells.

The source code will be made available at `https://github.com/Jacopo-Marchi/`.

### 2. PARAMETER ESTIMATION

In this section we detail the procedures we employed to fit parameters to reproduce our data or to qualitatively estimate them based on first principles, and we specify when we adopt parameters inferred in similar conditions in previous works. The resulting parameter values, biological meaning, and sources (when applicable) are reported in Table S1. When fitting a parameter to our experiments yielded a value comparable to a previously reported one, the "Source" column indicates the reference to the published value after the mention "Fitted". We stress here that most of the model parameters are effective parameters driving population dynamics in our phenomenological model. Whenever possible we constrained our parameter choices on studies inferring mechanistic quantities from microscale measurements such as [3–6, 10]. We clarify here that the qualitative dynamics presented in the main text are robust to variation in many of the fitted model parameters that do not find a counterpart in literature (see Section 2)– *i.e.* those without a citation on the "Source" column of Table S1. Therefore PhASER formation is not contingent on the details of the parameters estimation procedure presented in this Section. Instead, we used the parameters estimation procedure detailed below to capture quantitative features of our experimental measurements.

- Bacteria growth

  In order to fit bacteria life traits parameters, we consider experiments for each strain taking only into account phage-free portions of the swim assay, away from the lysis region. We take the radial intensity profile measured on a 0.25% agar plate inoculated with *E. coli* W1485(F8), far from phage T7 action (see Supplementary Fig. 21A), a proxy for bacteria density in the direction of range expansion. We normalize the intensity value to remove the artificial signal due to agar and scanner auto-calibration (Supplementary Fig. 21B). To do so we rescale the maximum intensity measured in the empty portion of the plate to the control intensity value measured in liquid medium without any bacteria in Supplementary Fig. 19. Then, we transform intensities to bacteria densities based on a spline interpolated on the intensity to CFU calibration data, as in Supplementary Fig. 19. First, we seek to infer the parameters related to bacterial growth, namely the initial amount of resources $R_0$, controlling the maximum bacterial density and the parameters controlling the bacterial growth profile, namely $K$ and $r$. The chemotactically migrating front seeds cells that start growing behind it [6]. We measure the speed $v$ at which the front is moving when the profile is recorded, allowing us to deduce that bacteria growing at a distance $\Delta$ behind the front have been seeded there at a previous time $t_s = \Delta/v$. Hence, the transformation $t = (r_{peak} - r)/v$ allows us to obtain the growth curve, *i.e.* bacteria densities as a function of time after seeding, from the spatial radial profile. We discard the part of the curve corresponding to the radial profile until the second chemotaxis front to avoid effects driven by bacteria transport rather than growth. We infer $R_0$, $r$ and $K$ by minimizing the squares

| Model parameters | | | | | |
|---|---|---|---|---|---|
| *Variable* | *Meaning* | *Value for different strains* | | *Units* | *Source* |
| | | *E. coli W1485*: | | | |
| | | (F8) | (F9) and (F10) (if different) | | |
| $r$ | Maximum bacteria growth rate | 1.3 | 1.6 | $\text{hrs}^{-1}$ | Fitted |
| $K$ | Growth Monod constant | $7 \cdot 10^9$ | $5 \cdot 10^9$ | CFU/mL | Fitted |
| $K_1$ | Aspartate Monod constant | 3.2 | 1.5 | $\mu$g/mL | Fitted |
| $K_2$ | Serine Monod constant | 10 | 3 | $\mu$g/mL | [1, 2], adjusted for mutants |
| $\mu_{1,\max}$ | Aspartate uptake rate | $2 \cdot 10^{-8}$ | $3 \cdot 10^{-8}$ | $\mu$g/(cell $\times$ hrs) | [3], adjusted for mutants |
| $\mu_{2,\max}$ | Serine uptake rate | $1.4 \cdot 10^{-7}$ | - | $\mu$g/(cell $\times$ hrs) | [3] |
| $a_-$ | Low chemosensing constant | 0.3 | - | $\mu$g/mL | [2, 4, 5] |
| $a_+$ | High chemosensing constant | 10 | 100 | $\mu$g/mL | Fitted, also [2, 6] |
| $D_B$ | Bacteria diffusion constant | $1.8 \cdot 10^5$ | - | $\mu\text{m}^2$/hrs | [6] |
| $\chi$ | Chemosensing responsiveness | $6 \cdot 10^6$ | $4 \cdot 10^6$ | $\mu\text{m}^2$/hrs | Fitted |
| $S_0$ | Initial cell density | $8 \cdot 10^8$ | - | CFU/mL | Experimental protocol |
| | | Phage: | | | |
| | | T7 | T4 | | |
| $\phi$ | Infection rate | $2 \cdot 10^{-8}$ | $6 \cdot 10^{-8}$ | mL/(hrs $\cdot$ PFU) | Fitted |
| $\eta$ | Maximum lysis rate | 5 | 2 | $\text{hrs}^{-1}$ | Fitted |
| $\beta$ | Burst size | 70 | 40 | PFU/CFU | Fitted |
| $\omega$ | Viral decay | 0.01 | - | $\text{hrs}^{-1}$ | [7] |
| $D_P$ | Phage diffusion constant | $10^4$ | - | $\mu\text{m}^2$/hrs | [8] |
| $L$ | Number of infected states | 10 | - | | [2, 9] |
| $V_0$ | Initial viral density | $10^9$ | - | PFU/mL | Experimental protocol |
| | | Chemicals properties: | | | |
| $D_R$ | Chemicals diffusion constant | $3 \cdot 10^6$ | - | $\mu\text{m}^2$/hrs | [10, 11] |
| $R_0$ | Initial resource level | $3.5 \cdot 10^9$ | - | CFU/mL | Fitted, also [2, 6] |
| $a_0$ | Initial chemoattractants level | 20 | - | $\mu$g/mL | [2, 6] |

TABLE S1: **Model parameters.** This table lists the baseline parameters used to simulate all strains. The values listed here for a given strain of bacteria or phage are used for all simulations featuring that strain, unless noted otherwise. The only parameters that are modulated from this baseline are $D_B$ and $\chi$ to reflect variations in range expansion speed given changes in the agar concentration, and the values reported here refer to the standard condition 0.25% agar. Notably, in order to avoid overfitting, phage infection parameters are consistently used across all simulations of different cell strains, although in general phage traits could depend on the host type if bacteria chemotaxis properties were pleiotropically linked to susceptibility. The procedure to fit or qualitatively estimate parameters, as well as the conversions between different units used in other works, is detailed below.

of the residuals between this data and a Monod growth model equivalent to the growth terms in Box 1, namely:

$$\frac{\partial R}{\partial t} = -r\frac{R}{R+K}S$$
$$\frac{\partial S}{\partial t} = r\frac{R}{R+K}S$$

where resources are expressed in units of cells, and bacteria start from a density $S_0$ fixed to the first data point in the curve. The optimization algorithm [12], applying the Levenberg-Marquardt method [13], yields $r = 1.1$ $\text{hr}^{-1}$ with an estimated standard error of 0.2 $\text{hr}^{-1}$, $K = 8 \pm 1.6 \cdot 10^9$ CFU/mL, and $R_0 = 2 \pm 0.1 \cdot 10^9$ CFU/mL to which we need to add the bacteria density after the second peak $S_0 = 1.6 \cdot 10^9$ to retrieve the corrected final bacteria density when starting from a small amount of cells, leading to $R_0 = 3.6 \pm 0.1 \cdot 10^9$ CFU/mL. To account for variability in our data we repeat this procedure on another 0.25% agar plate inoculated with *E. coli* W1485(F8), yielding $r = 1.3 \pm 0.7$ $\text{hr}^{-1}$, $K = 8 \pm 0.3 \cdot 10^9$ CFU/mL and $R_0 = 3.2 \pm 0.05 \cdot 10^9$ CFU/mL. We obtain estimates on the same order of magnitude, despite the fact that the preparation of the soft agar drives a certain degree of variability across experimental replicates. As final parameter estimates we select $R_0 = 3.5 \cdot 10^9$ CFU/mL, $K = 2R_0 = 7 \cdot 10^9$ CFU/mL and $r = 1.3$ $\text{hr}^{-1}$, compatible with the values inferred from both replicates. Supplementary Fig. 21C shows the growth curves from both plates, the growth model dynamics inferred on the first plate alone, and the growth curve produced by the final chosen parameters. Note that our value of $R_0$ is compatible with the final density of cells reported in previous range expansion studies in similar swim assays [2, 6].

We repeat the same procedure for *E. coli* mutants W1485(F9) and W1485(F10), estimating their parameters in the 0.3% agar plate where they first were sampled, see Supplementary Fig. 22A where the double-front *E. coli* W1485(F10) strain expands on the left of the lysis zone while the single-front *E. coli* W1485(F9) strain drives the dynamics on the right. Extracting the radial intensity profiles along the yellow segments (Supplementary Fig. 22) and then repeating the Monod growth parameters inference yields $r = 1.6 \pm 0.3$ hr$^{-1}$, $K = 5 \pm 0.08 \cdot 10^9$ CFU/mL and $R_0 = 2.5 \pm 0.03 \cdot 10^9$ CFU/mL for *E. coli* W1485(F9) and $r = 1.2 \pm 0.4$ hr$^{-1}$, $K = 5 \pm 0.08 \cdot 10^9$ CFU/mL and $R_0 = 2.3 \pm 0.04 \cdot 10^9$ CFU/mL for *E. coli* W1485(F10) (Supplementary Fig. 22C). As final parameters of the full model in Box 1 we choose both for *E. coli* W1485(F9) and *E. coli* W1485(F10) $r = 1.6 \pm 0.3$ hr$^{-1}$ and $K = 5 \pm 0.08 \cdot 10^9$ CFU/mL. In principle these two mutants can have different life traits, but we decided to use common parameters within a standard deviation of the inferred value to avoid overfitting. For the same reason, given variability in the swim assay preparation, we select a common $R_0 = 3.5 \cdot 10^9$ CFU/mL for all three bacteria strains.

Note that for brevity in the equations we rescaled resource densities in unit of CFU/mL, since scalar transformations between units do not impact the model dynamics provided that all parameters are rescaled accordingly. In some cases it can be useful to express all chemical concentrations, including resources, in $\mu$g/mL. This can be done based on a conversion factor indicating the amount of $\mu$g of resources constituting a cell, the yield $\epsilon$. *E. coli* cells typically contain $\sim$ 200 fg of carbon [14], therefore, if half of nutrient weight is due to carbon, it takes about 400 fg of processed nutrients to make up a cell. Assuming a cell metabolizes to growth slightly less than half of nutrients that are taken up, we estimate $\epsilon \approx 10^{-6}\mu$g/cell. In these units, dynamics are unchanged if the Monod constant, the initial resource concentration and the resource consumption rate (now different than bacteria growth rate) are multiplied by $\epsilon$.

We estimate the initial density of bacteria and phage inoculated from the stocks to be around $10^9$ per mL (see stocks section in Methods).

- Bacteria motility and sensing

We set the bacteria diffusion constant to $D_B = 1.8 \cdot 10^5 \mu\text{m}^2$/hrs [6]. Once $D_B$ is fixed, and with the maximum growth rate inferred as above, the sensing responsiveness parameter $\chi$ is inferred to match the range expansion speed, that at steady state scales as $v \propto \chi\sqrt{\frac{g(R_0)}{D_B}}$, as detailed in [6]. To reproduce the 5 mm/hr speed measured in 0.25% agar experiments we set $\chi = 6 \cdot 10^6 \mu\text{m}^2$/hrs. When extrapolating to other agar concentrations we change consistently both $D_B$ and $\chi$. To match the experiments at 0.2% agar in in Fig. 4A we obtain a speed of 8.3 mm/hr with $D_B = 3.2 \cdot 10^5$ and $\chi = 15.2 \cdot 10^6$, while $D_B = 10^5$ and $\chi = 2.9 \cdot 10^6$ yield a speed of 3.5 mm/h matching the experiments at 0.3% agar in Fig. 4B.

The low chemoattractant sensing constant was inferred in [4, 5] to be 3.5$\mu$M for the chemoattractant molecule aspartate, ($a_1$ in our model), which has $\approx$ 100g/mol = 0.1$\mu$g/($\mu$M$\cdot$mL), yielding approximately $a_- \approx 0.3\mu$g/mL. In the following we will use the same conversion factor from $\mu$M to $\mu$g/mL for serine, the other main chemoattractant in our swimming assays [2] ($a_2$ in our model), which has a similar molecular mass to aspartate. This conversion leads to a serine Monod constant of approximately $K_2 \approx 5 - 10\,\mu$g/mL [1, 2]. The uptake rate for serine $\mu_{2,\max}$ is 3.35 mmol/(g dry weight x hour) during cells exponential growth [3]. On average, a cell's dry mass amounts to $4 \cdot 10^{-13}$g [15], leading to a serine uptake of $3.4 \times 4 \cdot 10^{-13}$ mmol/(cell x hour), or $\mu_{2,\max} = 14 \cdot 10^{-8}\mu$g/(cell x hour). Applying the same conversion, we find an aspartate uptake rate per cell of $\mu_{1,\max} = 2 \cdot 10^{-8}\mu$g/(cell x hour) [3]. Finally, we estimate two additional life traits for the *E. coli* W1485(F8): the high sensing constant $a_+$ and the Aspartate Monod constant $K_1$, which we fit to reproduce the measured radial density profiles. We start from previously inferred $a_+ = 3\mu$g/mL [6] and $K_1 = 0.5\mu$g/mL [2], and we vary these values until the radial profile of a range expansion simulation resembles the data from the two reference plates used to fit the growth parameters (see Supplementary Fig. 21D). The final value for $a_+ = 10\mu$g/mL is on the same order of magnitude as the previously inferred parameter [6], and $K_1 = 3.2\mu$g/mL confirms a lower Monod constant for aspartate than for serine [2].

We repeat this procedure to fit chemosensing and motility parameters for *E. coli* mutants W1485(F9) and W1485(F10). In order to reproduce the experimental radial density profiles we set $a_+ = 100\mu$g/mL, which coincidentally corresponds to the value proposed in [2] if we apply the conversion $\approx$ 100g/mol introduced above, and $K_1 = 1.5\mu$g/mL. To reproduce the new mutants' profiles we also needed to adjust the serine Monod constant $K_2$ from 10 to 3$\mu$g/mL and the aspartate uptake rate $\mu_{1,\max}$ from $2 \cdot 10^{-8}$ to $3 \cdot 10^{-8}\mu$g/(cell x hour). This new set of chemosensing parameters yields the bacteria density profiles in Supplementary Fig. 22D, which closely reproduce the data. These radial curves are produced by simulations with $D_B = 10^5 \mu\text{m}^2$/hrs and

$\chi = 2.5 \cdot 10^6 \mu\text{m}^2/\text{hrs}$, as these values, similar to those used for *E. coli* W1485(F8) in Fig.4B, yield the $\sim$ 4mm/hr speed measured in the 0.3% agar plate in Supplementary Fig. 22A. To recover the $\sim$ 6.5mm/hr speed observed in 0.25% agar experiments for strain *E. coli* W1485(F10)(Fig.4C) and *E. coli* W1485(F9) (Fig.2B) we revert to $D_B = 1.8 \cdot 10^5 \mu\text{m}^2/\text{hrs}$ [6] and $\chi = 4 \cdot 10^6 \mu\text{m}^2/\text{hrs}$ – corresponding to the values reported in Table S1. Finally, to obtain the fast 7.3 mm/hr range expansion speed measured in Fig.2A we set $D_B = 2.7 \cdot 10^5 \mu\text{m}^2/\text{hrs}$ and $\chi = 5.5 \cdot 10^6 \mu\text{m}^2/\text{hrs}$. Other than the motility parameters to match the range expansion speeds measured in individual plates, given variations in the swim assay preparation, we assume for simplicity that strains *E. coli* W1485(F9) and *E. coli* W1485(F10) share all parameters, with *E. coli* W1485(F9) only responding to serine gradients (equivalent to $a_1 = 0$).

- Viral infection
  We select plausible values that are commonly used in phage-bacteria ecology studies to fix a few phage parameters, namely the viral decay $\omega$ from [7], phage diffusion constant $D_P = 10^4$ $\mu\text{m}^2/\text{hrs}$ from [8], and the number of infection steps leading to cell lysis $L = 10$ from [2, 9], which leads to a phage infection time variability in line with empirical estimates [16]. We vary the values of phage adsorption rate $\phi$, lysis rate $\eta$ and burst size $\beta$, selecting the values yielding the closest reproduction of the experimental dynamics, reported in Table S1, which qualitatively agree with previously reported phage infection parameters for T7 and T4 [7].

### 3. FITTED PARAMETER VALUES: COMPARISON WITH LITERATURE AND ROBUSTNESS ANALYSIS

Among the parameters estimated above, a subset does not find a quantitative match in literature in comparable experimental conditions – in these cases, a reference is not present in the "Source" column of Table S1. Here, we compare these parameters with previously published ones for transparency, while noting differences in experimental conditions. As mentioned at the beginning of Section 2, many of our parameters represent phenomenological terms driving population dynamics rather than mechanistic ground-truths. In this context, the quantitative agreement between our experiments and model justifies differences compared to previously published parameter values. To quantify the impact of such differences on the model dynamics we performed several robustness analyses, summarized here, varying fitted parameters. We organize the following using the same bullet points denoting different processes as in the previous parameters estimation Section.

- Bacteria growth
  The maximum bacteria growth rate $r$ was directly measured in nutrient-rich media between 1.1 and 1.4 $\text{hrs}^{-1}$ (and below 1.1 $\text{hrs}^{-1}$ in minimal media) [6], and was indirectly inferred as 2.8 $\text{hrs}^{-1}$ in [2] in soft agar experiments with LB medium. Our fitted $r$ values sit in between this range. Additionally, Supplementary Figure 14 C shows that T7 PhASER dynamics are robust to a 4-fold increase in growth rate, comfortably bracketing the value inferred in [2] (more details on the effects of modifying nutrients quality and assumptions are given in Section 4). The Monod constant $K$ was inferred in spatial agar assays with LB as about $10^9$ CFU/mL [2] (upon converting to our units). While our primary values are 5-7 times higher, simulations lowering $K$ to the same value of $10^9$ CFU/mL produce PhASERs both for T7 and T4 (Supplementary Figure 11 C and D respectively) with minor quantitative differences compared to our data.
- Bacteria motility and sensing
  The fitted aspartate Monod constants $K_1$, necessary to quantitatively reproduce our measured radial bacteria density profiles in Supplementary Figures 21 and 22, is on the same order of magnitude as inferred in LB soft agar assays [2]. As described above, once we fix all other bacteria parameters, the sensing responsiveness parameter $\chi$ is inferred to reproduce our measured range expansion speed. The values we inferred from experiments across conditions and strains range from $2.5 \cdot 10^6$ to $15.2 \cdot 10^6 \mu\text{m}^2/\text{hrs}$, producing the consistent formation of PhASERs at front speeds spanning 3.5 to 8.3 mm/hrs (see Figure 4 A,B). The lower end of this range is compatible with the $\chi \sim 1.5 \cdot 10^6 \mu\text{m}^2/\text{hrs}$ inferred in [2, 6], as are the front speeds measured in our experiments.
- Viral infection
  Moving to phage infection parameters, we note that typically phage life-history traits are quantified in liquid cultures. Therefore we cannot make direct comparisons for T7 and T4 phage parameters in the same environment as our experiments, where a number of factors ranging from spatial structure to nutrient dynamics could impact quantitative phage traits. With this caveat in mind, phage life-history traits were reported in [7] as $\phi = 3 \cdot 10^{-8}$ mL/(hrs · PFU), $\eta = 2.6$ $\text{hrs}^{-1}$, $\beta = 150$ PFU/CFU for phage T4, and $\phi = 1.8 \cdot 10^{-7}$ mL/(hrs · PFU), $\eta = 4.6$ $\text{hrs}^{-1}$, $\beta = 260$ PFU/CFU for phage T7. As mentioned above, different spatial settings could produce significant differences in the inferred $\phi$ value which implicitly accounts for diffusion/motility processes affecting phage-bacteria encounters, as $\phi$ scales linearly with phage diffusion constant [17]. Phage adsorption rate is also known

to significantly vary with cell surface in different growth conditions [18]. More broadly, phage life-history traits vary widely with nutrient availability and hosts growth rates [18, 19], a theme investigated in detail in Section 4. Modulating bacteria growth rates between 0.5 and 1.2 $\text{hrs}^{-1}$ approximate burst sizes were reported between 40 and 120 PFU/CFU for phage T7 (estimated from Fig. 2 of [19]). For phage T4, similar bacteria growth rates yield burst sizes between 20 and 60 PFU/CFU, and up to 110 PFU/CFU with a growth rate approaching 1.8 $\text{hrs}^{-1}$ [18].

Motivated by these large variations in measured phage life-history traits, we performed a robustness analysis varying phage infection parameters. First, we explored the simulated dynamics of phage transported by *E. coli* W1485(F9) when varying the magnitude of a linear adsorption rate and of lysis rate, spanning two orders of magnitude around the parameters inferred for phage T7. Supplementary Fig. 7 shows that for extremely low adsorption rates and lysis rates (corresponding to an average infection time of 1 hour) the phage infections are unable to sustain PhASERs. Increasing either adsorption or lysis rates (up to 4 minutes average infection period) produce stable PhASERs with a wider and wider lysis region, until the chemotactic fronts at its sides do not reconverge within our experiment's scales. Yet, within these two extreme limits, PhASER emergence shows a remarkable robustness to 10-fold variations in viral life traits, up to $\phi$ values close to those reported in [7]. Then we repeated the same procedure using the saturating adsorption profile of phage T4, spanning one order of magnitude around the corresponding $\eta$ and $\phi$ parameters (see Supplementary Figure 8). The realization of the observed experimental dynamics with residual cells inside narrow PhASERs appears to be more sensitive on these parameters values. High adsorption and lysis rates offset the nonlinear phage saturation profile and produce PhASERs that present T7-like features – i.e. initial curved fronts divergence and vanishing cell densities inside PhASERs. Finally, in Supplementary Figure 9 we increased the burst size $\beta$ both for phage T7 (panel A) and phage T4 (panel B), bracketing previously measured values [7, 18, 19]. With $\beta = 150$ PFU/CFU we recover qualitatively similar PhASER dynamics. A higher $\beta = 250$ PFU/CFU yields T7 lysis regions that do not reconverge within our experiment's scales, while T4 dynamics start presenting quantitative PhASER features typically associated with phage T7 as already noted for Supplementary Figure 8.

### 4. IMPACT OF CORE MODEL INGREDIENTS

Here we discuss in detail the remaining functional components constituting our mathematical model that were not addressed in Sections 2 and 3. To clarify the processes enabling the emergence of PhASERs, we tested the model behavior when modifying its core ingredients.

- **Phage discreteness threshold**
  The model relies on infection-mediated feedback determining the dependence of infection properties on resources and phage densities. We modulate all phage reaction terms by a nonlinear function $T(P) = \frac{P^M}{P^M + \Theta^M}$ with $M = 10$ to ensure that our continuous PDE model does not integrate early phage proliferation from nonphysical low phage densities. According to this 'single-phage form factor' phage rates sharply go to 0 when phage densities are lower than 1 virion per lattice site $P < \Theta = \frac{1}{V_l}$, with $V_l = 0.5 \cdot 10^{-4}$ being the volume of a site (see Section 1). Considering a simple well-mixed phage predation model with phage being washed out at rate $\alpha$ due to decay and spatial transport $\frac{\partial P}{\partial t} = (\beta - 1)\phi SPT(P) - \alpha P$, phage invading a phage-free system with $S^*$ bacteria grow ($\frac{\partial P}{\partial t} > 0$) if

$$P > \frac{\Theta}{\left(\frac{\phi(\beta-1)S^*}{\alpha} - 1\right)^{\frac{1}{M}}}.$$

  Therefore, with $M \gg 1$, phage can invade a dense bacteria lawn only above densities $P \gtrsim \Theta$, while with $T(P) = 1$ even a fraction of a virion would result in phage proliferation.

- **Non-exponential latent period distribution**
  Another key ingredient in our model is that infected cells progress through $L = 10$ infection stages between phage adsorption and lysis, corresponding to an Erlang distribution of phage infection times with a Coefficient of Variation (CV) of $1/\sqrt{10} \sim 0.3$ [9, 20]. To address the necessity of this ingredient we simulated the model dynamics with $L = 1$ [21], which corresponds to an exponentially distributed latent period, therefore increasing the CV to 1 while keeping the mean of the latent period distribution fixed. We found that an exponentially distributed infection time does not reproduce the experimental dynamics, as the high probability of lysis after a vanishing infection time produces a sideways expansion of the T7 lysis zone that outpaces the chemotactic

range expansion of adjacent uninfected bacteria. See Supplementary Fig. 4A showing a simulation where *E. coli* W1485(F8) confronts phage T7, with the same parameters as in Fig. 3 except for the latent period variance.

To characterize further the impact of lysis time distributions, we tested the effect of an increasing number of infected stages on $24cm$ plates dynamics simulated with the parameters otherwise inferred for either phage T7 or T4 (Supplementary Figure 5). Again, assuming an exponentially distributed latent period leads to a lysis zone that keeps expanding outwards for more than 30 cm, diverging drastically from our empirical observations, as shown in Supplementary Figure 5, panel A for T7 and D for T4. Increasing the number of infected stages produces a narrower unimodal latent period distribution, with a smaller Coefficient of Variation (decreasing as $1/\sqrt{L}$) around the fixed average lysis time. This yields a faster reconvergence of the initial lysis zone, although with $L = 2$ we observe an instability of T7 PhASERs to fronts merging (panel B). This outcome is impacted by phage life history traits, since T4 parameters with $L = 2$ produce a stable PhASER (panel E). Introducing 3 infected stages was sufficient to produce a long-range PhASER regardless of phage parameters (panel C for T7 and F for T4), therefore the dynamics stabilize well ahead of the $L = 10$ value employed in the main text. This finding is consistent with empirical measurements of the latent period's CV, which are well below 1 and are compatible our choice of 10 infected stages [16]. Note that smaller $9cm$-diameter plates (dotted blue circle in Supplementary Figure 5) do not permit sufficient space to assess the curvature of initial lysis boundaries [21] making it hard to distinguish between exponential and peaked distributions, and consequently prevent the characterization of PhASERs stability.

- **Coupling between resources and phage infection**

In our model the transition rate from phage adsorption to lysis is coupled to cell metabolism through resource concentration as $\eta(R) = \eta \frac{R}{R+K}$ as is the burst size $\beta(R) = 1 + \beta \frac{R}{R+K}$, which are therefore linearly related to bacteria growth rate. Supplementary Fig. 4B shows a simulation with *E. coli* W1485(F8) *vs* T7 where phage infections do not slow down as resources are depleted, with $\eta(R) = \eta/3$ ensuring the same basal lysis rate as in Fig. 3 of the main text since $K = 2R_0$. After phage infections are transported by the range expansion front (gray overlay showing 2h-spaced fronts), phage slowly consume the bacterial lawn through a constant expansion of the lysis region behind the front.

Then, we sought to test the qualitative impact of introducing a nonlinear dependence of lysis rate and burst size on bacteria growth rate. A quantitative characterization of this dependence in spatial environments where bacteria dynamically run out of nutrients is currently not available. Therefore we turned our attention to measurements of the dependence of phage life traits with hosts growth rates in well-mixed cultures where bacteria grow at steady state. While noting the different conditions with respect to our experiments, which could impact the quantitative growth-dependent phage traits functions, this choice allows us to qualitatively test nonlinear functions informed by empirical observations. Specifically, we leverage the work presented in [22], where the authors inferred the dependence of phage life traits with host growth rate from previous experiments on phage T7 [19] and T4 [18, 23]. An empirically-informed evolutionary steady state analysis in [22] yields the latent period (reported in minutes therein) as a function of host growth rate $\tilde{r}$ normalized by the maximum growth rate measured for each phage $r$, as:

$$l(r_{\text{norm}}) = \frac{1}{w} + E_\infty + E_0\, e^{-\alpha_E r_{\text{norm}}},$$

and a burst size given by:

$$\beta(r_{\text{norm}}) = \frac{M_\infty}{w} \cdot \frac{1}{1 + e^{-\alpha_M (r_{\text{norm}} - M_0)}},$$

with $r_{\text{norm}} = \frac{\tilde{r}}{r}$. In our model, $r_{\text{norm}} = \frac{R}{R+K}$, which decreases as resources $R$ are consumed. The parameters in these function were inferred in [22] from [19] for phage T7 as $E_\infty = 15.33$ min, $E_0 = 82.71$ min, $\alpha_E = 5.9$, $M_\infty = 6.7$ PFU/(CFU $\cdot$ min), $M_0 = 0.64$, $\alpha_M = 9.7$, and from [18, 23] for phage T4 as $E_\infty = 18.26$ min, $E_0 = 94.18$ min, $\alpha_E = 7.76$, $M_\infty = 47.32$ PFU/(CFU $\cdot$ min), $M_0 = 0.65$, $\alpha_M = 12.03$. These functions depend on a free parameter $w$ , which we set so that the initial lysis rate, with resources at the initial concentration $R_0$ (Table S1), is the same as in the main text simulations. In other words, we set $w$ so that $l\left(r_{\text{norm}} = \frac{R_0}{R_0+K}\right)/60 = \left(\eta \frac{R_0}{R_0+K}\right)^{-1}$, where the term 60 rescales the time units from minutes to hours, consistently with our units. This choice allows us to test the impact of adding this nonlinearity, without affecting the range of the lysis rates realized in our simulations based on the resource concentration dynamics. This leads to a nonlinear growth-dependent lysis rate:

$$\eta(r_{\text{norm}}) = \frac{\eta \frac{R_0}{R_0+K}}{1 + \frac{\eta}{60} \frac{R_0}{R_0+K} E_0 \left(e^{-\alpha_E r_{\text{norm}}} - e^{-\alpha_E \frac{R_0}{R_0+K}}\right)}, \tag{S1}$$

where the constant $\eta$ in the right hand side denotes the parameter specified in Table S1. Similarly, to consistently renormalize the burst size to the correct initial value, we set $w$ so that $\beta\left(r_{\mathrm{norm}}=\frac{R_0}{R_0+K}\right)=1+\beta\frac{R_0}{R_0+K}$. This yields a nonlinear burst size:

$$\beta(r_{\mathrm{norm}})=\left(1+\beta\frac{R_0}{R_0+K}\right)\frac{1+e^{-\alpha_M(\frac{R_0}{R_0+K}-M_0)}}{1+e^{-\alpha_M(r_{\mathrm{norm}}-M_0)}}, \tag{S2}$$

where the right hand side constant $\beta$ is specified in Table S1.

Supplementary Figure 10 shows the results of spatial simulations employing these nonlinear phage traits functions. Panels A and B respectively show eq. (S1) and eq. (S2) (full lines), compared to the linear function employed in the main text (dashed lines), both for phage T7 (red) and T4 (blue). The x-axis spans the normalized growth rate range realized in the simulations based on the resources dynamics parameters – *i.e.* $[0, \frac{R_0}{R_0+K}]$. Despite the quantitative functional differences, these nonlinear phage traits produce PhASER dynamics very similar to the linear traits presented in Figure 2 in the main text (panel C for phage T7 and panel D for phage T4).

To test the impact of nonlinear lysis rate and burst size even further, we decrease the Monod constant $K$ from $5\cdot10^9$ CFU/mL to $10^9$ CFU/mL, almost doubling $\frac{R_0}{R_0+K}$ and consequently the range of normalized growth rates and phage traits spanned in the spatial simulations. This choice exacerbates the nonlinearities in lysis rate and burst size in equations (S1) and (S2) (see Supplementary Figure 11 A and B respectively). First, we run simulations with this modified parameter using the original linear dependence (dashed lines in Supplementary Figure 11 A and B) of lysis rate and burst size with growth rate. The spatial simulations results are shown in Supplementary Figure 11 C for phage T7 and in Supplementary Figure 11 D for phage T4. As already mentioned in the previous Section, the 5-fold reduction in $K$ produces the typical qualitative PhASER dynamics as in the main text both for phage T7 and T4. Then, we run new simulations using the nonlinear dependence expressed in equations (S1) and (S2) (full lines in Supplementary Figure 11 A and B), both for phage T7 (Supplementary Figure 11 E) and T4 (Supplementary Figure 11 F). The nonlinearities in lysis rate and burst size, more pronounced in these conditions compared to those simulated in Supplementary Figure 10, still do not affect the emergence of stable PhASERs, despite small quantitative differences in the simulated dynamics.

- **Nonlinear T4 phage adsorption rate and phage infection strength**

  The simulations reproducing phage T4 experimental large-scale dynamics presented in the main text employ an adsorption profile saturation at high phage densities $F(P)=\phi\frac{P}{1+P/P_c}$ with $P_C=10^7$ PFU/mL [20, 24], following Michaelis-Menten reaction kinetics [25] which mediate inter-phage competition processes such as lysis inhibition [26]. This saturation profile helps capture more features of T4 PhASERs, such as the presence of a low density of cells coexisting with phage within the lysis region. Yet, we highlight here that we can reproduce most of the qualitative features of T4 PhASERs with a simpler linear adsorption $F(P)=\phi P$ similar to simulated T7 dynamics. Indeed, Supplementary Fig. 17 shows T4-like simulated dynamics resulting from a linear adsorption profile with $\phi=2\cdot10^{-9}$ mL/(hrs $\cdot$ PFU), $\eta=5$ hrs$^{-1}$ and $\beta=50$ PFU/cell, while the rest of the parameters are fixed to the *E. coli* W1485(F8) bacterium strain values reported in Table S1. These ingredients yield a narrow T4-like PhASER propagated over long distances (Supplementary Fig. 17) which transfers to coalescing colonies expanding at different speeds provided that the angle of incidence between the merging fronts is not too big (Supplementary Fig. 17), as in the infection transport dynamics presented in Fig. 5 in the main text.

- **Chemically distinct nutrients and attractants**

  We tested the model predictions merging nutrients and attractants into the same state variable [21], ignoring the chemical complexity of the LB medium. To do so, we rescaled the parameters setting the concentration band of maximum sensitivity, $a_-$ and $a_+$, multiplying them by the ratio of carrying capacities $K/K_2$ to match the order of magnitude of resource dynamics, and we set $\chi=5\cdot10^7\mu\mathrm{m}^2/\mathrm{hrs}$ to keep the range expansion speed comparable to experiments. Supplementary Figure 6 shows that conflating nutrients and attractants leads to qualitatively different phage propagation dynamics. Keeping the rest of the parameters unchanged with respect to those inferred for T7, including the average later period ($\eta=5$/hr) (panels A,B,C), we see that bacteria wash away all phage and invade the whole plate, due to the fact that the bacteria population saturates to carrying capacity and limits lysis right behind the front (see [6]). At intermediate time snapshots before bacteria fill the infected sector (A,B) the boundaries of the phage-bacteria interface appear to be straight as reported by [21]. This outcome is even clearer when we increase the lysis rate $\eta$ just enough to stop bacteria invasion at the center of the plate, setting $\eta=10$/hr (D,E,F), which leads to straight boundaries throughout the simulated dynamics.

- **Biomass conversion and nutrients quality**

  To complement our experimental investigation on the impact of environmental variations (Figure 4), we extended the numerical exploration of the model addressing its robustness to nutrients quality, another environment feature, focusing on *E. coli* strain W1485(F9). To do so, we vary the biomass conversion factor $\epsilon$ (yield) between nutrients measured in $\mu$g/mL and cells measured in CFU/mL, introduced in Section 2. As noted when we discussed our parametrization of bacterial growth, choosing a different unit of measure for nutrients, expressed in CFU/mL in the main text, does not affect the model dynamics.

Setting $\epsilon = 5 \cdot 10^{-7} \mu$g/cell halves the yield with respect to the original parameters (lower nutrient quality or metabolic efficiency, corresponding to a slower initial doubling time of 2.6 hours) and the T7 phage are transported with a widened initial lysis region (Supplementary Figure 14 A). Supplementary Figure 14 B reproduces the main T7 simulations with $\epsilon = 10^{-6} \mu$g/cell for comparison. When the yield is set to 4 times the original parameter cells grow faster, corresponding to a 20 minutes initial doubling time, and a PhASER emerges with a narrower lysis region (Supplementary Figure 14 C). Note that as predicted by [6], changing the growth rate affects the chemotactic range expansion speed. The high yield conditions in panel C produce a fast 15 mm/hr range expansion, which is comparable to chemotactic front speeds reported in liquid media [27]. This result suggests that the model qualitative dynamics are robust to nutrient quality perturbations.

---

[7] De Paepe, Marianne; Taddei F (2006) Viruses' life history: Towards a mechanistic basis of a trade-off between survival and reproduction among phages. *PLOS Biology* 4:null.

[8] Barr JJ, et al. (2015) Subdiffusive motion of bacteriophage in mucosal surfaces increases the frequency of bacterial encounters. *Proceedings of the National Academy of Sciences* 112:13675–13680.

[9] Mitarai N, Brown S, Sneppen K (2016) Population dynamics of phage and bacteria in spatially structured habitats using phage λ and escherichia coli. *Journal of Bacteriology* 198:1783–1793.

[10] Cremer J, et al. (2016) Effect of flow and peristaltic mixing on bacterial growth in a gut-like channel. *Proceedings of the National Academy of Sciences* 113:11414–11419.

[11] Fraebel DT, et al. (2017) Environment determines evolutionary trajectory in a constrained phenotypic space. *eLife* 6:e24669.

[12] Newville M, Ingargiola A, Stensitzki T, Allen DB (2014) *Lmfit: Non-Linear Least-Square Minimization and Curve-Fitting for Python.*

[13] Moré JJ (1978) *The Levenberg-Marquardt algorithm: Implementation and theory* ed Watson GA (Springer Berlin Heidelberg, Berlin, Heidelberg), pp 105–116.

[14] J.S. W (2015) *Quantitative Viral Ecology: Dynamics of Viruses and Their Microbial Hosts*, Monographs in Population Biology (Princeton University Press, Princeton), p 360.

[15] Neidhardt FC, Curtiss R, eds (1996) *Escherichia coli and Salmonella: Cellular and Molecular Biology* (ASM Press, Washington, D.C.), 2nd edition.
